# *In vivo* CRISPRa decreases seizures and rescues cognitive deficits in a rodent model of epilepsy

**DOI:** 10.1101/431015

**Authors:** Gaia Colasante, Yichen Qiu, Luca Massimino, Claudia Di Berardino, Jonathan H. Cornford, Albert Snowball, Mikail Weston, Steffan P. Jones, Serena Giannelli, Andreas Lieb, Stephanie Schorge, Dimitri M. Kullmann, Vania Broccoli, Gabriele Lignani

## Abstract

Epilepsy is a major health burden, calling for new mechanistic and therapeutic insights. CRISPR–mediated gene editing shows promise to cure genetic pathologies, although hitherto it has mostly been applied *ex-vivo*. Its translational potential for treating non-genetic pathologies is still unexplored. Furthermore, neurological diseases represent an important challenge for the application of CRISPR, because of the need in many cases to manipulate gene function of neurons *in situ*. A variant of CRISPR, CRISPRa, offers the possibility to modulate the expression of endogenous genes by directly targeting their promoters. We asked if this strategy can effectively treat acquired focal epilepsy, focusing on ion channels because their manipulation is known be effective in changing network hyperactivity and hypersynchronisation. We applied a doxycycline-inducible CRISPRa technology to increase the expression of the potassium channel gene *Kcna1* (encoding Kv1.1) in mouse hippocampal excitatory neurons. CRISPRa-mediated Kv1.1 upregulation led to a substantial decrease in neuronal excitability. Continuous video-EEG telemetry showed that AAV9-mediated delivery of CRISPRa, upon doxycycline administration, decreased spontaneous generalized tonic-clonic seizures in a model of temporal lobe epilepsy, and rescued cognitive impairment and transcriptomic alterations associated with chronic epilepsy. The focal treatment minimizes concerns about off-target effects in other organs and brain areas. This study provides the proof of principle for a translational CRISPR-based approach to treat neurological diseases characterized by abnormal circuit excitability.

## Introduction

Epilepsy affects up to 1% of the population, and 30% of patients continue to experience seizures despite the use of current medication (Kwan *et al*., 2011; Tang *et al*., 2017). Although the majority of drug-resistant epilepsies are focal, targeting drugs to a restricted brain region presents major challenges, and potentially curative surgery is limited to a minority of cases where the seizure focus is remote from eloquent cortex (Kullmann *et al*., 2014). Gene therapy holds promise as a rational replacement for surgery for intractable pharmaco-resistant epilepsy, and could in principle improve the prospect for seizure freedom in many people (Kullmann *et al*., 2014; Lieb *et al*., 2018). Several approaches have been proposed to interfere with epileptogenesis or to decrease seizure frequency in chronic epilepsy (Simonato, 2014). Current experimental gene therapies mainly rely on viral vector-mediated expression of genes encoding normal CNS proteins or exogenous non-mammalian proteins (Wykes *et al*., 2012; Krook-Magnuson *et al*., 2013; Katzel *et al*., 2014; Wykes *et al*., 2016; Lieb *et al*., 2018). This approach has several potential limitations, including a finite packaging capacity of viral vectors, difficulty in ensuring normal splicing and post-transcriptional processing, and, in the case of non-mammalian proteins, concerns about immunogenicity. Modulating the expression of endogenous genes, in contrast, would represent an important step toward safe and rational treatment of intractable epilepsy and other neurological diseases.

The DNA editor/regulator CRISPR/Cas9 (Konermann *et al*., 2015; Dominguez *et al*., 2016; Adli, 2018) represents a powerful tool to modify endogenous genes, not only in somatic cells but also in mammalian neurons (Heidenreich and Zhang, 2016; Suzuki *et al*., 2016). In addition to permanently altering endogenous gene sequences, CRISPR/Cas9 can regulate the activity of genes through promoter modulation, an approach known as CRISPR activation (CRISPRa) (Dominguez *et al*., 2016; Liao *et al*., 2017). CRISPRa is therefore a promising tuneable tool to increase the expression of genes encoding, for instance, ion channels, in chronic epilepsy in order to restore physiological levels of network activity (Wykes *et al*., 2012; Wykes and Lignani, 2018). The CRISPRa system consists of a nuclease-defective Cas9 (dCas9) fused to a transcription activator and a small guide RNA (sgRNA) that targets dCas9 to the promoter of the gene of interest (Dominguez *et al*., 2016). There are multiple advantages of this system. First, it is versatile because the targeted gene can be switched simply by changing the sgRNA. Second, CRISPRa preserves the full range of native splice variants and protein biogenesis mechanisms (Liao *et al*., 2017). Third, CRISPRa is, in principle, safe because it only alters the promoter activity of genes that are already transcribed in targeted neurons. Finally, CRISPRa can be targeted to specific neurons in the epileptic focus using established viral vectors (La Russa and Qi, 2015).

Here, we report the use of CRISPRa to treat a mouse model of temporal lobe epilepsy, from *in vitro* validation to demonstration of efficacy in reducing seizure frequency and rescuing cognitive impairment *in vivo*.

## Materials and Methods

### Study Design

This study aimed to test the hypothesis that upregulating endogenous genes (e.g. *Kcna1*) with CRISPRa can treat chemoconvulsant-induced temporal lobe epilepsy. The experiments were designed to achieve a power >0.8 with an α = 0.05. For *in vivo* experiments the 3Rs guidelines for animal welfare were followed. Outliers were not excluded and at least 3 independent repetitions were performed. Exclusion criteria were applied for all the recordings (see methods below). All the experiments were randomized and researchers were blinded during recordings and analysis.

### Animals and ethics

All experimental procedures were carried out in accordance with the UK Animals (Scientific Procedures) Act 1986. C57BL/6J and Camk2a-CRE mice were used for the experiments. Animals have been housed in groups or single after surgery in IVC cage in an SPF facility.

### Plasmids

sgRNAs were cloned in a lentiviral pU6 vector. Ef1alpha-dCas9VP160-T2A-PuroR, was derived from pAC94-pmax-dCas9VP160-2A-puro, a gift of R. Jaenisch (Addgene plasmid # 48226). The dCas9VP160-2A-puro cassette was subcloned in a TetO-FUW vector and then restriction digested with HpaI/AfeI and blunt cloned into an Ef1alpha-GFP vector after GFP removal by SmaI/EcoRV digestion. Ef1alpha-dCas9VP160-T2A-GFP was obtained by restriction digestion of Ef1alpha-dCas9VP160-T2A-PuroR with AscI/XbaI that removed VP160-T2A-PuroR; the VP160-T2A fragment was then obtained by AscI/XhoI digestion from Ef1alpha-dCas9VP160-T2A-PuroR while GFP fragment was PCR amplified with primers containing XhoI/XbaI restritction sites; finally, the two fragments were ligated together into the vector. To obtain a single vector containing both dCas9A and sgRNA, the pU6-sgRNA cassette was HpaI digested and cloned into Ef1alpha-dCas9VP160. AAV-TRE-dCas9-VP64 was obtained by restriction digestion of AAV-SpCas9 (gift of F. Zhang, Addgene # PX551) where the Mecp2 promoter was removed by XbaI/AgeI digestion and TRE promoter was amplified with the following primers: FWXbaI: GCTCTAGACCAGTTTGGTTAGATCTC and RV AgeI GCACCGGTGCGATCTGACGGTTCACT. SpCas9 was removed with AgeI/EcoRI and Cas9m4-VP64 (gift of G. Church, addgene # 47319) was digested with AgeI/EcoRI. The VP64 fragment was PCR amplified with following primers with EcoRI sites: F: GATCATCGAGCAAATAAGCGAATTCTC and R: gctaaGAATTCTTA-TCTAGAGTTAATCAGCATG. The AAV vector containing the sgRNA cassette was derived from pAAV-U6sgRNA (SapI)_hSyn-GFP-KASH-bGH (PX552 was a gift of F. Zhang, Addgene # 60958): sg19 or lacZ where cloned under U6 promoter and the GFP was removed by KpnI/ClaI digestion and replaced by a DIO-rtTA-T2A-Tomato cassette. This vector was used for the work in Camk2a-Cre mice. For the work in C57/Bl6 mice, this vector was XbaI/ClaI-digested to remove both hSyn promoter, and the DIO-rtTA-T2A-Tomato cassette was replaced by a Camk2a promoter amplified with NheI-KpnI and a tTA T2a tomato cassette amplified with KpnI ClaI, ligated together in the vector.

### Virus preparation

Lentiviruses were produced as previously described with a titer of 10^7-10^8 IU/ml (Colasante *et al*., 2015). AAVs were produced as previously described with a titer higher than 10^12 vg/ml (Morabito *et al*., 2017). The TRE-dCas9-VP64 AAV was produced by VectorBuilder with a titer of 8 × 10^12 vg/ml.

### P19 cell line

P19 cells were cultured in alpha-MEM (Sigma-Aldrich) supplemented fetal bovine serum non-essential amino acids, sodium pyruvate, glutamine and penicillin/streptomycin and split every 2-3days using 0.25% trypsin. For transfection, Lipofectamine 3000 (Thermo Fisher Scientific) was used according to the manufacturer’s instructions.

### Primary neuronal culture and lentivirus transduction

Cortical neurons were isolated from P0 C57Bl/6J mouse pups as previously described (Beaudoin *et al*., 2012) and at 1 DIV were transduced with lentiviruses. qRT-PCR, RNA seq, Western blot analysis and electrophysiology recordings were performed 14-16 days after transduction.

### RNA isolation and quantitative RT-PCR

RNA was extracted using TRI Reagent (Sigma) according to the manufacturer’s instructions. For quantitative RT-PCR (RT-qPCR), cDNA synthesis was obtained using the ImProm-II Reverse Transcription System (Promega) and RT-qPCR were performed with custom designed oligos (Table 1) using the Titan HotTaq EvaGreen qPCR Mix (BIOATLAS). Analysis of relative expression was performed using the ΔΔC_T_ method, relative to Ctrl-dCas9A condition. To determine Kcna1 expression in vivo, ΔΔC_T_ was determined in Ctrl-dCas9A or in Kcna1-dCas9A injected hippocampi relative to contralateral hippocampi in epileptic animals at the end of the recordings.

### Western Blot

Total neuronal protein extracts were obtained from the lysis of primary neurons by RIPA lysis buffer (150 mM NaCl, 1% Triton, 0.5% sodium deoxycholate, 0.1% sodium dodecyl sulfate, Tris pH 8.0 50 mM, protease inhibitor cocktail) two weeks after infection with the CRISPRa-Kcna1 system. Lysates were kept on ice for 30 minutes by vortexing every 10 minutes and then centrifuged at 4° C for 5 minutes at 5000 rpm. Supernatants with solubilized proteins were collected in new tubes and stored at -80° C until use. Western blot analysis was performed using primary antibodies against the following proteins: anti-Kv1.1 (1:1000, Neuromab) anti-α/βActin (1:10000, Sigma).

### Bioinformatic analysis

Encyclopedia of DNA Elements (ENCODE) and the Functional ANnoTation Of the Mammalian genome (FANTOM) (Carninci *et al*., 2006) databases were used to download transcriptomics and epigenetics NGS data. Tracks were visualized along the mm10 mouse reference genome with the Integrative Genome Viewer (IGV) (Thorvaldsdottir *et al*., 2013).

### Off-targets

Employing the free Galaxy web-tool (https://usegalaxy.org/) we generated two datasets: one containing sg19 off-target sequences predicted by *CRISPOR* web tool (http://crispor.tefor.net) and one containing all the 500 bp genomic regions (NCBI37/mm9) upstream to transcription start sites (TSS) of annotated transcripts. Intersecting the two datasets, all sg19 off-target sequences in putative gene promoters were derived. To identify genes regulated by putative promoters, the sequence of the predicted off-targets was aligned by IGV to the reference genome and to transcripts annotated in ENSEMBL database. Validation of expression levels of putative off-target genes was performed by RT-qPCR.

#### RNA-seq

RNA libraries for both in vitro and in vivo experiments were generated starting from 1 ug of total RNA extracted from sglacz- and sg19-dCas9A neurons at 10 DIV. RNA quality was assessed by using a Tape Station instrument (Agilent) and only RNA samples with Integrity Number (RIN) ≥ 8 were analyzed. For in vitro experiment, RNA was processed according to the Lexogen QuantSeq 3′ mRNA-Seq Library Prep Kit protocol and the libraries were sequenced on an Illumina NextSeq 500 with 75bp stranded reads at CTGB, Ospedale San Raffaele. Fastq files were aligned to the mouse genome (NCBI37/mm9) with Bowtie2.

For in vivo experiments, RNA was processed according to the TruSeq Stranded mRNA Library Prep Kit protocol. The libraries were sequenced on an Illumina HiSeq 3000 with 76 bp stranded reads using Illumina TruSeq technology at Genewiz. Image processing and base calling were performed using the Illumina Real Time Analysis Software. Fastq files were mapped to the mm10 mouse reference genome with the STAR aligner v2.7 (Dobin *et al*., 2013).

Differential gene expression and functional enrichment analyses were performed with DESeq2 (Love *et al*., 2014) and GSEA, respectively. Statistical and downstream bioinformatics analyses were performed within the R environment. Gene expression heatmaps were produced with GENE-E (Broad Institute). Data of in vitro and in vivo experiments were deposited in the NCBI Gene Expression Omnibus repository with the GSE133930 GEO ID.

### Slice preparation

Camk2a-CRE mice of either sex (2-3 months old) were killed by cervical dislocation under isoflurane. Brains were quickly dissected into ice cold oxygenated slicing solution (in mM: 75 sucrose, 2.5 KCl, 25 NaHCO_3_, 25 glucose, 7 MgCl_2_, 0.5 CaCl_2_) and cut into 300 µm coronal slices using a Leica VT1200S vibratome (Leica). Slices were stored submerged in oxygenated recording artificial cerebrospinal fluid (aCSF) (in mM: 25 glucose, 125 NaCl, 2.5 KCl, 25 NaHCO_3_, 1 MgCl_2_, 1.25 NaH_2_PO_4_.H_2_O and 2 CaCl_2_) at 32°C for 30min and at room temperature for a further 30min before recording.

### Electrophysiology

*In vitro.* For current-clamp recordings, the internal solution contained (in mM): 126 K-gluconate, 4 NaCl, 1 MgSO4, 0.02 CaCl2, 0.1 BAPTA, 15 Glucose, 5 HEPES, 3 ATP-Na2, 0.1 GTP-Na, pH 7.3. The extracellular (bath) solution contained (in mM): 2 CaCl2, 140 NaCl, 1 MgCl2, 10 HEPES, 4 KCl, 10 glucose, pH 7.3. D-(−)-2-amino-5-phosphonopentanoic acid (D-AP5; 50 μM), 6-cyano-7-nitroquinoxaline-2,3-dione (CNQX; 10 μM) and picrotoxin (PTX; 30 μM) were added to block synaptic transmission. Transduced excitatory neurons were identified with EGFP fluorescence and from a pyramidal somatic shape. Neurons with unstable resting potential (or > -50mV), access resistance (R_a_) > 15 MΩ and/or holding current >200 pA at -70mV were discarded. Bridge balance compensation was applied and the resting membrane potential was held at -70 mV. A current step protocol was used to evoke action potentials (APs), by injecting 250 ms long depolarizing current steps of increasing amplitude from -20pA (Δ 10pA). Recordings were acquired using a Multiclamp 700A amplifier (Axon Instruments, Molecular Devices) and a Power3 1401 (CED) interface combined with Signal software (CED), filtered at 10 kHz and digitized at 50 kHz.

#### Ex-vivo current clamp recordings

Current clamp recordings were performed in standard external solution in the presence of DL-AP5 (50 μM), 6-cyano-7-nitroquinoxaline-2,3-dione (CNQX; 10 μM) and PTX (30 μM) to block NMDA, AMPA/kainate, and GABA_A_ receptors, respectively. The internal solution was the same as for in vitro patch clamp recordings. Neurons with holding current >100 pA and R_a_ >20 MΩ upon whole-cell break-in in voltage clamp mode and membrane potential less negative than -60mV in current clamp were not considered for analysis. A 1440 Digidata (Molecular Devices) or Power3 1401 (CED) interface and Multiclamp 700A (Molecular Devices) amplifier were used.

#### In vitro and ex-vivo electrophysiology analysis

All the electrophysiology analysis was performed with an automated Python script. Passive properties were calculated from the hyperpolarizing steps of the current clamp steps protocol. Input resistance was averaged from three current steps (2 negative and one positive). Capacitance was calculated from the hyperpolarizing current step as follows. Firstly, the input resistance was determined as the steady-state ΔV/ΔI (voltage/current), then the cell time constant (tau) was obtained by fitting the voltage relaxation between the baseline and the hyperpolarizing plateau. Capacitance was then calculated as tau/resistance. Single action potential parameters were calculated as previously described (Pozzi *et al*., 2013). An event was detected as an action potential if it crossed 0mV and if the rising slope was >20mV/ms in a range of injected currents from 0pA to 500pA. All the experiments were performed at room temperature (22-24°C). All recordings and analysis were carried out blind to vector transduced.

#### Activity clamp

The template simulating the barrage of synaptic conductances during epileptiform bursts was previously described (Morris *et al*., 2017). Dynamic clamp software (Signal 6.0, Cambridge Electronic Design, Cambridge, UK) and a Power3 1401 (CED) were used to inject both excitatory and inhibitory conductance templates simultaneously in a neuron recorded in current clamp configuration (iteration frequency 15 kHz). E_rev_ was set to 0 mV and -75 mV for excitatory and inhibitory conductances respectively, and corrected for a liquid junction potential of 14.9 mV. Incrementing synaptic conductances were injected in recorded neurons to establish the conductance threshold for action potential generation. Current clamp recordings for activity clamp were performed with the same external and internal solutions as given above.

### Surgical procedures

All the surgery procedures were performed in adult mice (2-3 months) anesthetized and placed in a stereotaxic frame (Kopf).

#### Epileptic model

0.3μg of 10mg/ml kainic acid (Tocris) was injected in a volume of in 200nl in the right amygdala (AP: -0.94; ML: 2.85; DV: 3.75) at 200nl/min under isoflurane anaesthesia (surgery time 10-15 minutes). The mice were allowed to recover from anaesthesia at 32°C for 5 minutes and then moved back to their cage where they were monitored closely during status epilepticus (SE). SE usually started 10-15 minutes after complete recovery and always stopped 40 minutes after KA injection with 10mg/Kg intraperitoneal diazepam.

#### Stereotaxic viral injection

300nl of AAV9 viruses (1:1) were injected with a 5μl Hamilton syringe (33 gauge) at 100nl/min in 3 different coordinates of the right ventral hippocampus (Antero-Posterior: -2/3 bregma/lamda distance; Medio-Lateral: -3; Dorso-Ventral: 3.5/3/2.5). the needle was kept in place for 10 minutes after each injection.

#### Transmitter implantation

An electrocorticogram (ECoG) transmitter (A3028C-CC Open Source Instruments, Inc) was subcutaneously implanted and the recording electrode was placed above the viral injection site (Antero-Posterior: -2/3 bregma/lamda distance; Medio-Lateral: - 3). The ground electrode was placed in the contralateral frontal hemisphere.

#### Exclusion criteria

Only animals recorded for the entire period of the experiment (6 weeks after KA) were used in the analysis. At the end of the experiments some animal tissues were analysed with qRT-PCR and others were verified with immunofluorescence. On total of 42 mice injected with kainic acid, 34 animals (80%) were implanted and injected. 24 were recorded for entire duration of the experiment. 2 did not express dCas9 and for this reason were excluded from the analysis. 22 mice were used for the analysis (13 Ctrl-dCas9A and 9 Kcna1-dCas9A).

#### Pilocarpine acute seizure model

Male wild type C57BLC/6J mice (3 months old) were anaesthetized with isoflurane and placed in a stereotaxic frame (David Kopf Instruments Ltd., USA). The animals were injected with 1.5μl AAV CamKII-CRISPR-Kcna1/CamKII-CRISPR-LacZ at 100nl/min in layer 2/3 - 5 primary visual cortex (coordinates: AP -2.8mm, ML 2.4 from the bregma, and DV 0.7/0.5/0.3 from pia). For EcoG monitoring, the recording electrode of 256Hz single-channel EcoG transmitter (A3028C-CC, Open Source Instruments Inc., USA) was placed at the same coordinates. A reference electrode was placed in the contralateral skull. A cannula (Bilaney Consultants Ltd., UK) was implanted in the same location as the recording electrode for sequential pilocarpine injections. Animals were allowed to recover for 2 weeks before induction of acute seizures by pilocarpine (3.5M in saline) (Magloire *et al*., 2019) injected 0.5mm below the cannula using a microinjection pump (WPI Ltd., USA), a 5 µl Hamilton syringe (Esslab Ltd., UK), and a 33 gauge needle (Esslab Ltd., UK). The injection volume was incremented on consecutive days (180nl, 300nl and 500nl) until spike-wave discharges were observed, and recorded as the threshold dose. If seizures failed to terminate spontaneously, the animal was excluded from the study. To assess the treatment, the animals were placed on a doxycycline diet for 7 days and only the threshold dose for the animal was repeated. EcoG monitoring was used to assess seizure severity for an hour after the pilocarpine injection. The researcher who acquired and analysed the data was blinded to the virus injected.

### EEG (or ECoG) recordings

The ECoG was acquired wirelessly using hardware and software from Open Source Instruments, Inc. The ECoG was sampled at a frequency of 256Hz, band-pass filtered between 1 and 160Hz, and recorded continuously for the duration of the experiments. The animals were housed independently in a Faraday cage.

### EEG analysis

Spontaneous seizures were detected from chronic recordings using a semi-automated supervised learning approach (suppl. figure 7). First, a library containing examples of epileptiform activity was built using seizures identified from visual inspection of ECoG data. The recordings were saved in hour-long files, and for each seizure this full hour was included in the library. Recordings were chunked into 5 second blocks that were labelled as either “ictal” or “interictal” if they contained epileptiform-labelled activity or not, respectively. For each five second chunk of recording, 15 features were extracted (suppl. figure 7 and see online resource below). A random forest discriminative classifier was trained on the features and labels of each of the 5 second examples in the library (Breiman, 2001). In addition, cross validation generated classifier predictions were used to parameterise a Hidden Markov Model in which the hidden states were the human annotations and the emissions the classifier predictions. For automated detection of epileptiform activity from unlabelled recordings, the discriminative classifier was first used to predict the class of consecutive five second chunks. We then applied the forward-backward algorithm to obtain the marginal probability of being in seizure state for each recording chunk given the surrounding classifier predictions. The smoothed predictions were then manually verified, false positives removed from the analysis and start and end locations adjusted. In order to quantify the performance of our approach, we randomly selected four 2 week chunks of recordings and visually examined the traces for seizures and compared to classifier predictions (blinded). During the 8 weeks, we did not detect visually any seizures that were not marked by the classifier – as such, for this model of epilepsy, our false negative rate was less than 1/300. False positives were less of a concern, but in general we observed << n seizures for a given period of time. For further information and code, please see: https://github.com/jcornford/pyecog.

### Video recordings

IP cameras from Microseven (https://www.microseven.com/index.html) were used and synchronised via the Windows time server to the same machine as used to acquire the ECoG. Continuous video recordings produced 6 videos/hour.

### Immunohistochemistry

Immunostaining was performed on 50µm mice brain sections with the following antibodies: mouse anti-GAD67 (MAB5406, Merck), rabbit anti-RFP (600-401-379, Rockland), Alexa Fluor 555 goat anti-rabbit (A32732, Invitrogen) and Alexa Fluor 488 goat anti-mouse (A32723, Invitrogen). Images were acquired with ZEN software (Zeiss) on a LSM710 confocal microscope (Zeiss) and co-localization analysis of tdTomato and GAD67 were performed with ImageJ 1.51n (Wayne Rasband, National Institute of Health) plugin ‘JACoP’.

#### Behavior Tests

Trials started two weeks post-virus injection, all were carried out between 7 a.m. - 7 p.m. during the light phase. Animals (3 months old) were habituated in the designated behavior room for at least 15 minutes in home cages prior to the test.

##### Object Location Test (OLT)

For the familiarization phase, mice were placed individually in the arena (50 cm x 50cm x 40 cm), and were allowed to explore two identical objects for 8 mins placed in the arena at least 5 cm away from the border. After a 6-hour retention delay, the animals were returned to the same arena with one of the objects randomly relocated to a new location. The animal was allowed to explore for 8 minutes with video recordings. The arena and objects were thoroughly cleaned with ethanol between each session.

##### Novel Object Recognition Test (NORT)

24 hours after the OLT, the same animals were subjected to the NORT test. The familiarization session was the same as for the OLT. After a 6 hours retention delay, one of the objects was randomly replaced by a novel object with a different shape and surface texture. The animals were allowed to explore freely for 8 minutes (Leger *et al*., 2013).

All trials were recorded with a Raspberry Pi 3B+ equipped with a V1 camera module (https://www.raspberrypi.org/documentation/hardware/camera/) and using Raspivid version 1.3.12 as 1296×972 pixel, 30 frame/second MP4 video files. Automated analysis was carried out with custom scripts written in Bonsai version 2.4-preview (Lopes *et al*., 2015). A researcher blinded to the treatment assessed and scored the exploration time manually after automated analysis. Discrimination index (DI) was calculated using the following formula: (time spent with altered object - time spent with unchanged object)/(total time spent exploring objects).

### Statistical Analysis

Data are plotted as box and whiskers, representing interquartile range (box), median (horizontal line), and max and min (whiskers), together with all the points. The mean is further shown as “+”. The statistical analysis performed is shown in each figure legend. Deviation from normal distributions was assessed using D’Agostino-Pearson’s test, and the F-test was used to compare variances between two sample groups. Student’s two-tailed *t-*test (parametric) or the Mann-Whitney test (non-parametric) were used as appropriate to compare means and medians. Fisher’s exact test was used to analyze the contingency table. To compare two groups at different time points we used two-way repeated measure ANOVA, followed by Bonferroni post-hoc test for functional analysis. Statistical analysis was carried out using Prism (GraphPad Software, Inc., CA, USA) and SPSS (IBM SPSS statistics, NY, USA).

### Data availability

All the Python and Bonsai scripts are freely available. Plasmids will be deposited in Addgene, and transcriptomic data are deposited in the NCBI Gene Expression Omnibus repository with the GSE133930 GEO ID.

## Results

### A CRISPRa system targeting the Kcna1 promoter region increases Kv1.1 expression and decreases neuronal excitability

We first asked if CRISPRa can be exploited to increase endogenous gene expression in glutamatergic neurons and decrease their excitability. As a proof of principle, we chose the *Kcna1* gene encoding the Kv1.1 channel, which is important for the regulation of neuronal action potential firing and synaptic transmission (Pinatel *et al*., 2017; Vivekananda *et al*., 2017). Lentivirus- or adeno-associated virus-mediated overexpression of *Kcna1* reduces neuronal excitability and, when targeted to principal cells, suppresses seizures in rodent models of epilepsy (Wykes *et al*., 2012; Snowball *et al*., 2019). We first conducted a bioinformatic analysis to identify its promoter region. Alignment of datasets of gene expression and epigenetic markers of actively transcribed genes in perinatal and adult mouse brain identified peaks of enrichment for RNA PolII, mono- and tri-methylation of lys4 and acetylation of lys27 of H3 histone along the gene (Supplementary Figure 1A). One of these regions was located immediately upstream to the annotated *Kcna1* transcription start site (TSS) and identified as a suitable target for CRISPRa. We submitted 200 bps from this region to the *CRISPOR* web tool (http://crispor.tefor.net) for sgRNA design, and selected four candidate guides (sg4, sg14, sg19 and sg30) for validation, initially in the mouse P19 cell line that expresses many neuronal genes. sgRNAs were lipofected individually or in combination, together with a construct carrying dCas9 fused to the transcriptional activator VP160 (dCas9-VP160) and a Puromycin resistance cassette. dCas9 with sg4, sg14 or sg19, but not with sg30, significantly upregulated the expression of the *Kcna1* gene. We focused on sg4 and sg19, which induced the highest levels of *Kcna1* expression (Supplementary Figure 1B). When tested in combination, sg4 and sg19 together were also efficacious, but not sg4 and sg30 (Supplementary Figure 1C). We confirmed that the highest efficiency of upregulation of *Kcna1* in primary neurons was achieved with sg19 (Figure 1B). Consequently, we generated a construct carrying dCas9-VP160 driven by the Ef1 promoter and either the sg19 targeting the *Kcna1* promoter (Kcna1-dCas9A) or a control sgRNA targeting LacZ (Ctrl-dCas9A). Western Blot analysis confirmed increased Kv1.1 protein levels in sg19-treated neurons when compared to the sgLacZ control. Importantly, we detected increased levels of glycosylated Kv1.1, corresponding to mature protein, implying normal processing of the upregulated potassium channel (Figure 1C, D) (Watanabe *et al*., 2003).

**Figure 1:**
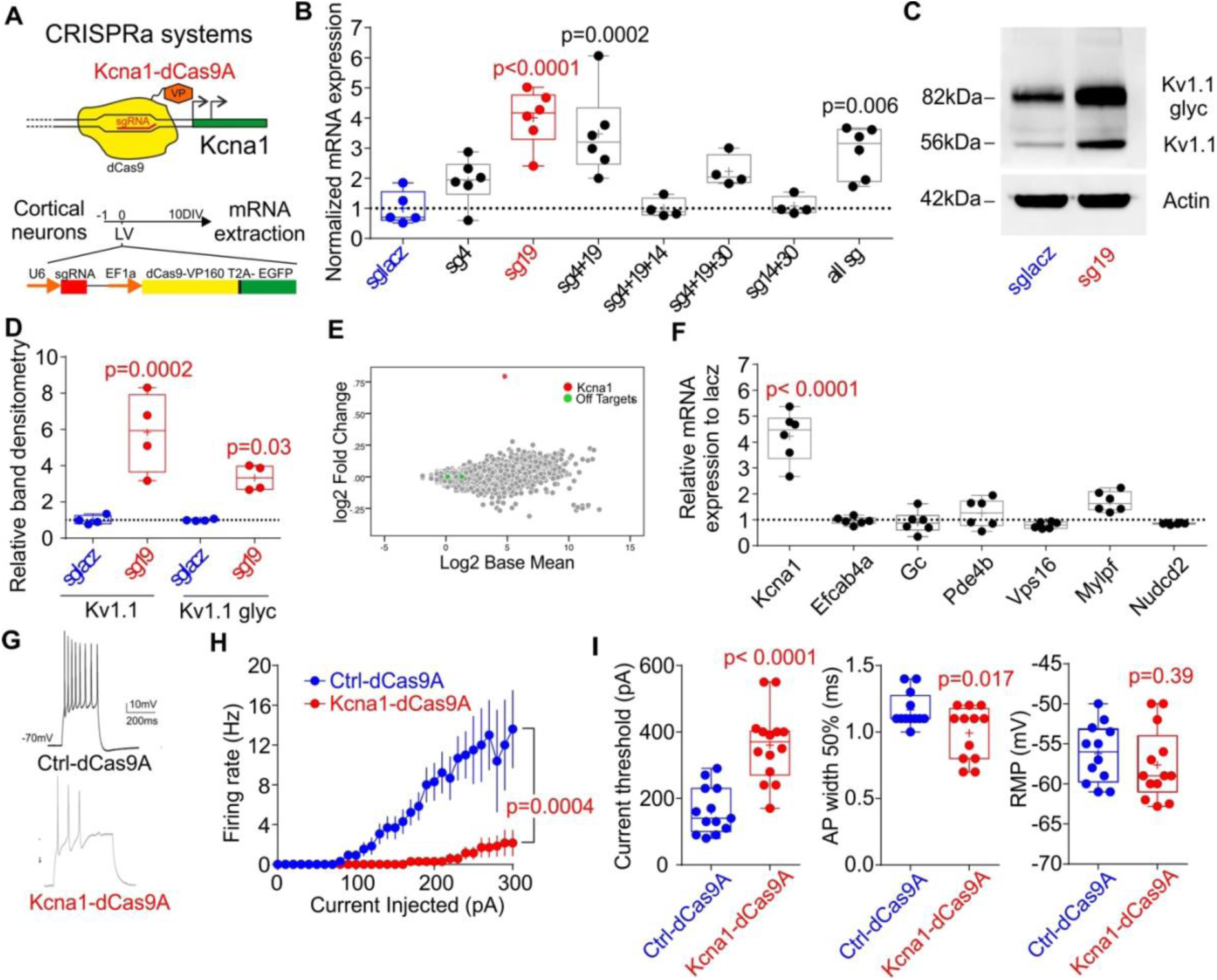
CRISPRa increases endogenous *Kcna1* expression reducing neuronal excitability *in vitro*. **A.** Schematic representation of the CRISPRa approach to increasing *Kcna1* **B.** *Kcna1* mRNA expression normalised to the control LacZ sgRNA (blue) for the most effective sgRNAs and combinations of different sgRNAs tested in P19 cells (Suppl. Figure 1). One-way ANOVA followed by Bonferroni multiple comparison test vs sgLacZ. **C-D.** Western blots were used to determine the increase in Kv1.1 and glycosylated Kv1.1 (glyc) in neurons transduced either with dCas9A and sg19 (red), or sgLacZ (blue). Student’s t test. **E.** MA plots showing log2 Fold change as a function of log2 base mean expression of *Kcna1*-dCas9A treated neurons with respect to Ctrl-dCas9A. *Kcna1*, red dot; off-targets, green dots; all other genes, gray dots. **F.** mRNA expression relative to sgLacZ for each off target. The expression level of sgLacZ is represented as the dashed line at 1. Multiple Student’s t tests, each compared to control and corrected for multiple comparison (α= 0.0083). **G.** Representative traces of recordings from neurons transduced either with Ctrl-dCas9A (sgLacZ, blue) or Kcna1-dCas9A (sg19, red) and injected with 300pA steps in current clamp. **H.** Average firing rates in response to different current injections for neurons transduced with ctrl-dCas9A or Kcna1-dCas9A. Two-way RM ANOVA. **I.** Neuronal and action potential (AP) properties in neurons transduced with ctrl-dCas9A or Kcna1-dCas9A. Student’s t test.

The *CRISPOR* tool predicted 250 putative off-target genes for sg19, mostly with a very low likelihood score. To evaluate the specificity of CRISPRa we performed a gene expression profile analysis in primary neurons treated with Kcna1-dCas9A and compared this with Ctrl-dCas9A transduced neurons. No consistent alteration in the transcriptome of sg19 treated neurons was observed, except for a significant increase in *Kcna1* (red dot, Figure 1E). Six out of 250 predicted off-target genes for sg19 were located close to promoters of *Mylpf, Efcab4a, Nudcd2, Pde4b, Gc* and *Vps16* genes. However, none of these genes showed a significant change in expression in either the transcriptome analysis (green dot, Figure 1E) or in quantitative RT-PCR assays (Figure 1F).

Exogenous *Kcna1* overexpression results in a decrease in neuronal excitability and neurotransmitter release (Heeroma *et al*., 2009; Wykes *et al*., 2012). To test the functional efficacy of the CRISPRa system, primary neurons were transduced at 1DIV with a lentivirus expressing Kcna1-dCas9A or Ctrl-dCas9A. After 14-16DIV we used whole-cell patch clamp recordings to analyse neuronal excitability of both experimental groups (Figure 1G). The maximal firing frequency was significantly decreased in neurons transduced with Kcna1-dCas9A when compared to Ctrl-dCas9A (Figure 1H). Other excitability parameters sensitive to Kv1.1 were also changed in neurons transduced with Kcna1-dCas9A in comparison with Ctrl-dCas9A: the current threshold was increased, and action potential width was decreased (Figure 1I). Passive membrane properties and other AP properties were however unchanged (Supplementary Figure 2).

### Kcna1-dCas9A decreases CA1 pyramidal cell excitability

In order to test the efficacy of CRISPRa *in vivo*, we subcloned the CRISPRa elements in two separate AAV9 vectors. One AAV vector carried the dCas9-VP64 under the control of a rtTA responsive element (TRE), while the other vector included the sg19 (or sgLacZ as a control) element and a human Synapsin promoter (hSyn) upstream to an inverted rtTA-t2a-tdTomato cassette flanked by loxP and lox511 sites. This experimental design allowed the Kcna1-dCas9A system to be activated in forebrain principal neurons of Camk2a-cre mice transduced with both AAVs, but only after doxycycline administration (Figure 2A, B). We co-injected both AAVs in the hippocampus of 2-3 month old Camk2a-CRE mice, which were subsequently fed with a doxycycline diet for 3 weeks before collecting acute brain slices for analysis. Pyramidal neurons in the CA1-subiculum of the ventral hippocampus were recorded with whole-cell patch clamp to measure their excitability (Figure 2B, C and Supplementary Figure 3, 4). Consistent with data from primary cultures, neurons transduced with Kcna1-dCas9A showed a decreased firing rate and increased current threshold when compared with Ctrl-dCas9A expressing neurons (Figure 2E). A minor difference from primary cortical cultures was that the half width of the first spike was not significantly different between Kcna1-dCas9A and Ctrl-dCas9A expressing neurons (Figure 1I). However, a significant decrease in half width was seen when all the APs during the current step protocol were pooled (Figure 2 F). Finally, we applied activity clamp, a method to assess neuronal excitability during epileptiform barrages of excitation (Morris *et al*., 2017). Neurons expressing Kcna1-dCas9A fired less than neurons expressing Ctrl-dCas9A when exposed to the same simulated synaptic input. Taken together, these results support using Kcna1-dCas9A as a candidate antiepileptic gene therapy (Figure 2 G).

**Figure 2:**
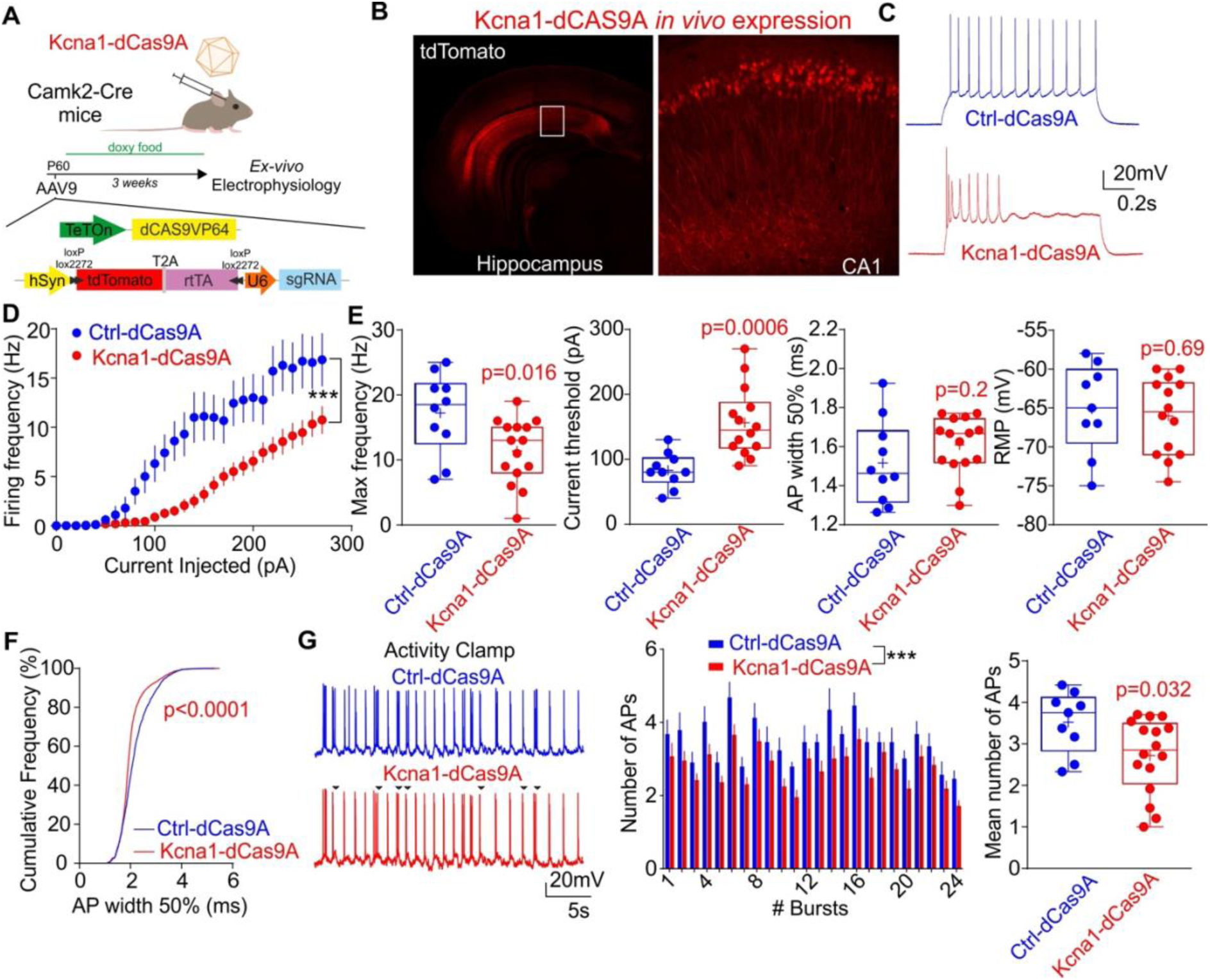
CRISPRa delivered with AAV9 increases endogenous *Kcna1* expression and reduces CA1 pyramidal neuron excitability. **A.** Schematic representation of the approach for *ex vivo* quantification of CRISPRa effects. **B.** Representative live image of *Kcna1-*dCas9A expression in the hippocampus (inset= CA1 region, Red= tdTomato) **C.** Representative traces from CA1 pyramidal neurons, transduced either with Ctrl-dCas9A (sgLacZ, blue) or Kcna1-dCas9A (sg19, red) in pryramidal neurons injected with 280pA steps in current clamp. **D.** Average firing rates in response to different current steps for neurons transduced with Ctrl-dCas9A or Kcna1-dCas9A. Two-way RM ANOVA. **E.** Neuronal and action potential (AP) properties in neurons transduced with Ctrl-dCas9A or Kcna1-dCas9A. Maximal firing rate, current threshold to elicit the first AP, AP width and resting membrane potential are shown. Student’s t test. **F.** Cumulative frequency (%) of the AP widths in neurons injected with 280pA of current. Kolmogorov-Smirnov test for cumulative distributions. **G.** Activity clamp protocol to mimic 24 interictal bursts of synaptic input from an epileptic network in neurons transduced with Ctrl-dCas9A or Kcna1-dCas9A (*left*). Black arrows represent the APs missed in the Kcna1-dCas9A neurons. Number of APs for each burst showed as mean ± sem (*middle*). Two-way ANOVA. The histogram represents the average number of APs for each neuron in the 24 bursts (*right*). Student’s t test.

### Kcna1-dCas9A decreases seizure frequency in a mouse model of temporal lobe epilepsy

We administered Kcna1-dCas9A in a mouse model of acquired epilepsy. C57BL/6J wild type animals were injected with kainic acid (KA) in the right amygdala (Jimenez-Mateos *et al*., 2012). This induced a period of status epilepticus (SE), which was quantified by video recording to monitor seizure severity (Supplementary Figure 5 and Video 1). One week later, we injected either Kcna1-dCas9A or Ctrl-dCas9A AAVs in the right ventral hippocampus, and at the same time we implanted wireless EEG transmitters (bandwidth 1-256 Hz). For these experiments an AAV9 carrying a rtTA-t2a-tdTomato cassette without flanking recombination sites but driven by a Camk2a promoter was delivered in order to bias expression to excitatory neurons. After a week of recovery to allow expression of the constructs, we started continuous video-EEG recording for 2 weeks (baseline) and then continued recording for 2 further weeks with doxycycline administration (Figure 3A). Immunohistochemistry of the injected hippocampi showed expression of the tdTomato reporter in dentate gyrus granule cells and hippocampal CA3 excitatory neurons, as well as CA1 pyramidal cells (Figure 3B, Supplementary Figure 6). We extracted both the ipsilateral and contralateral hippocampi of 11 mice after the EEG recordings to analyse *Kcna1* and dCas9 expression. Injected hippocampi from 2 mice failed to express dCas9 and were excluded from the analysis. The injected hippocampi from the remaining 9 mice (5 Ctrl-dCas9A and 4 Kcna1-dCas9A) expressed dCas9 and showed a 50% increase in *Kcna1* expression in mice transduced with Kcna1-dCas9A compared to controlateral counterpart while no difference is reported in Ctrl-dCas9A (Figure 3C).

**Figure 3:**
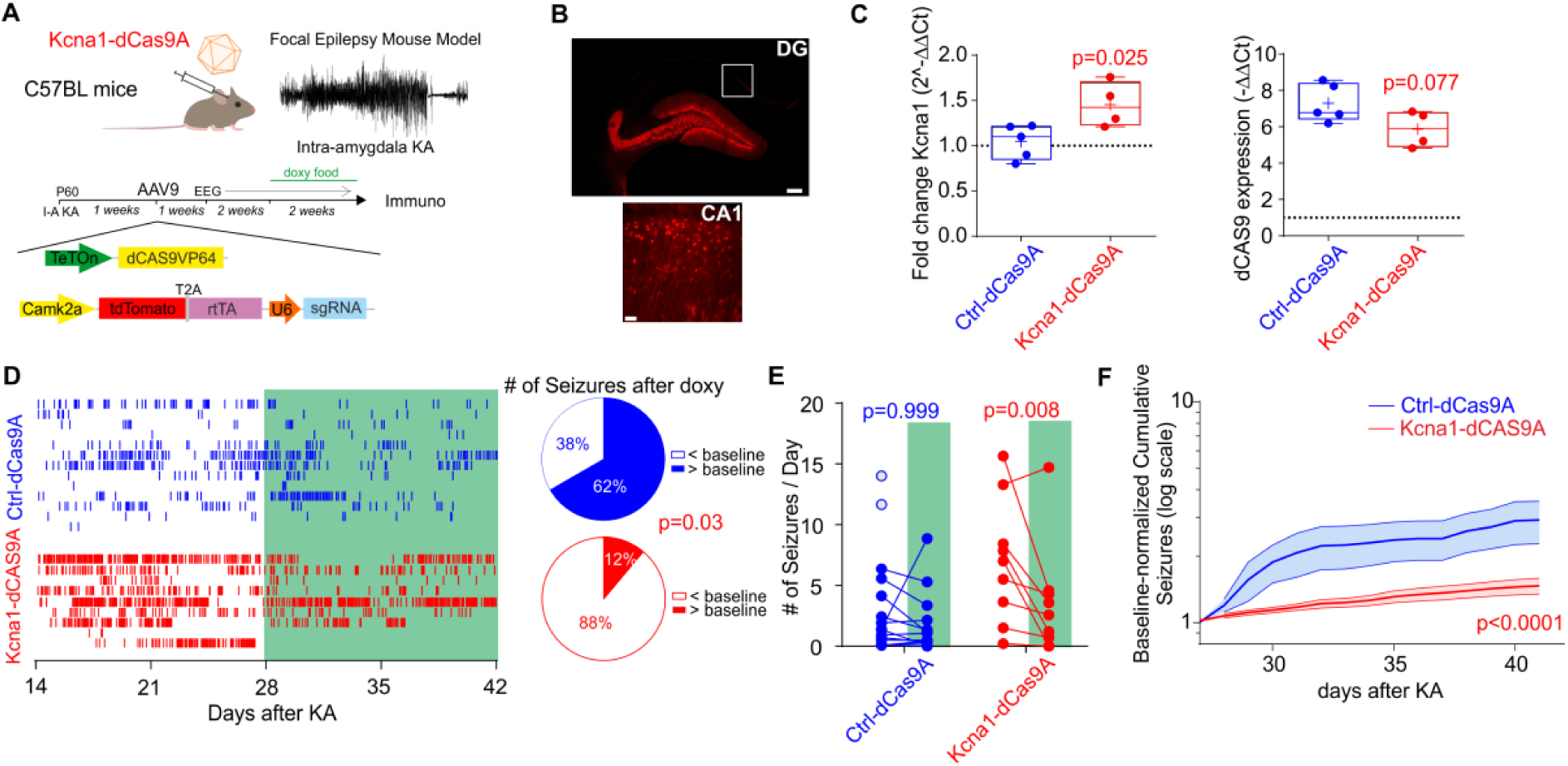
CRISPRa-Kcna1 decreases number of seizures in a mouse model of acquired intractable temporal lobe epilepsy. **A.** Schematic representation of the CRISPRa approach used *in vivo* to treat the intra-amygdala kainic acid focal model of temporal lobe epilepsy. **B.** Representative immunofluorescence 7 weeks after status epilepticus (SE) of neurons transduced with Ctrl-dCas9A 4 weeks after SE. Scale bar DG: 250 m; CA1:50 m. **C.** qPCR analysis of Kcna1 and dCas9 expression in the ipsilateral hippocampus relative to the contralateral hippocampus in epileptic mice transduced with either Ctrl-dCas9A or Kcna1-dCas9A at the end of the experiments. Student’s t test. **D.** *Left.* Raster plot showing all seizures before and after doxycycline administration in 13 mice treated with Ctrl-dCas9A and 9 mice treated with Kcna1-dCas9A. *Right.* Pie charts showing the proportion of animals showing either more or fewer seizures after doxycycline food than during the baseline. Fisher’s exact test. **E.** Number of seizures/day before and after doxycycline administration in control-dCas9 (n=13) and Kcna1-dCas9 (n=9) treated animals. Two-way ANOVA followed by Bonferroni multiple comparison test. Empty blue circles are animals that died during baseline and excluded from the analysis. **F.** Cumulative plot of seizures normalised to the baseline in mice transduced with either ctrl-dCas9A or Kcna1-dCas9A. Two-way ANOVA.

To investigate the ability of Kcna1-dCas9 to treat chronic temporal lobe epilepsy we analysed the frequency of generalized tonic-clonic seizures (Racine stage 5) in each animal before and after doxycycline administration using continuous video-EEG recordings (Figure 3D-F; Supplementary Figure 7; Video 2). Animals treated with Kcna1-dCas9A showed a significant reduction in the number of seizures after doxycycline was added to the food (Figure 3D-F). The number of seizures after doxycycline administration decreased in 8 out of 9 animals treated with Kcna1-dCas9A compared to only 5 out of 13 animals treated with Ctrl-dCas9 (Figure 3D). Only Kcna1-dCas9A treated animals showed a significant decrease in the number of seizures per day after doxycycline administration (Figure 3E, F). There was no significant difference in seizure frequencies during the baseline period prior to doxycycline administration between the animals treated either with Kcna1-dCas9A or Ctrl-dCas9A (Figure 3E, p=0.09 two-way ANOVA followed by Bonferroni multiple comparison test). However, two Ctrl-dCas9A animals, but none of the Kcna1-dCas9A animals, died during the baseline period from severe seizures and were not included in the comparison of seizure frequencies before and after doxycycline (Figure 3E). We cannot exclude a protective effect of a low level of basal activation of CRISPRa in the Kcna1-dCas9A animals. Other EEG parameters such as broadband power, seizure duration and EEG power for night-time and day-time periods were not changed by the treatment (Supplementary Figure 5 and 7). Taken together, these results show that Kcna1-dCas9A reduces the probability of tonic-clonic seizure initiation, but otherwise does not change seizure properties recorded in the cortex.

We complemented the chronic epilepsy study by looking at the effect of Kcna1-dCas9A treatment on acute seizures evoked by a different mechanism, focal pilocarpine injection in the visual cortex before and after doxycycline administration (Lieb *et al*., 2018; Magloire *et al*., 2019). We did not observe robust differences in EEG coastline and power in different frequency bands recorded for 1 hour after pilocarpine in either group, although there was a non-significant trend for Kcna1-dCas9 animals to show an attenuation in seizure severity (Supplementary Figure 8). These data argue that the action of Kcna1-dCas9A on experimental epilepsy goes beyond an effect on individual seizures.

### Kcna1-dCas9A rescues cognitive deficits and mitigates dysregulation of gene expression

Cognitive co-morbidities are an important feature of many forms of intractable epilepsy. We therefore asked if Kcna1-dCas9A treatment rescues behavioural deficits in our chronic intra-amygdala KA model of temporal lobe epilepsy. We used two behavioural paradigms, one directly related to hippocampal function (object location test, OLT) and the other more related to prefrontal cortex function (novel object recognition test, NORT) as previously described (Cho *et al*., 2015). In the OLT test, no impairment in performance, as measured by the discrimination index, before and after doxycycline, were observed in non-epileptic (NE) animals treated with either Ctrl-dCas9A or Kcna1-dCas9A (Figure 4A-C). However epileptic (E) mice showed a deficit compared to control mice (Ctrl-dCas9A NE vs E, p=0.01; Kcna1-dCas9A NE vs E, p=0.006; two-way ANOVA followed by Bonferroni multiple comparison test). This deficit was completely rescued only by Kcna1-dCas9A treatment (Figure 4C). In contrast, no significant differences were observed in the NORT test either between NE and E animals, or following treatment with either Ctrl-dCas9A or Kcna1-dCas9A (Supplementary Figure 9) (Cho *et al*., 2015). These data suggest that while Kcna1-dCas9A does not have adverse effects on hippocampal function in non-epileptic mice, it is able to restore function in epileptic mice.

**Figure 4:**
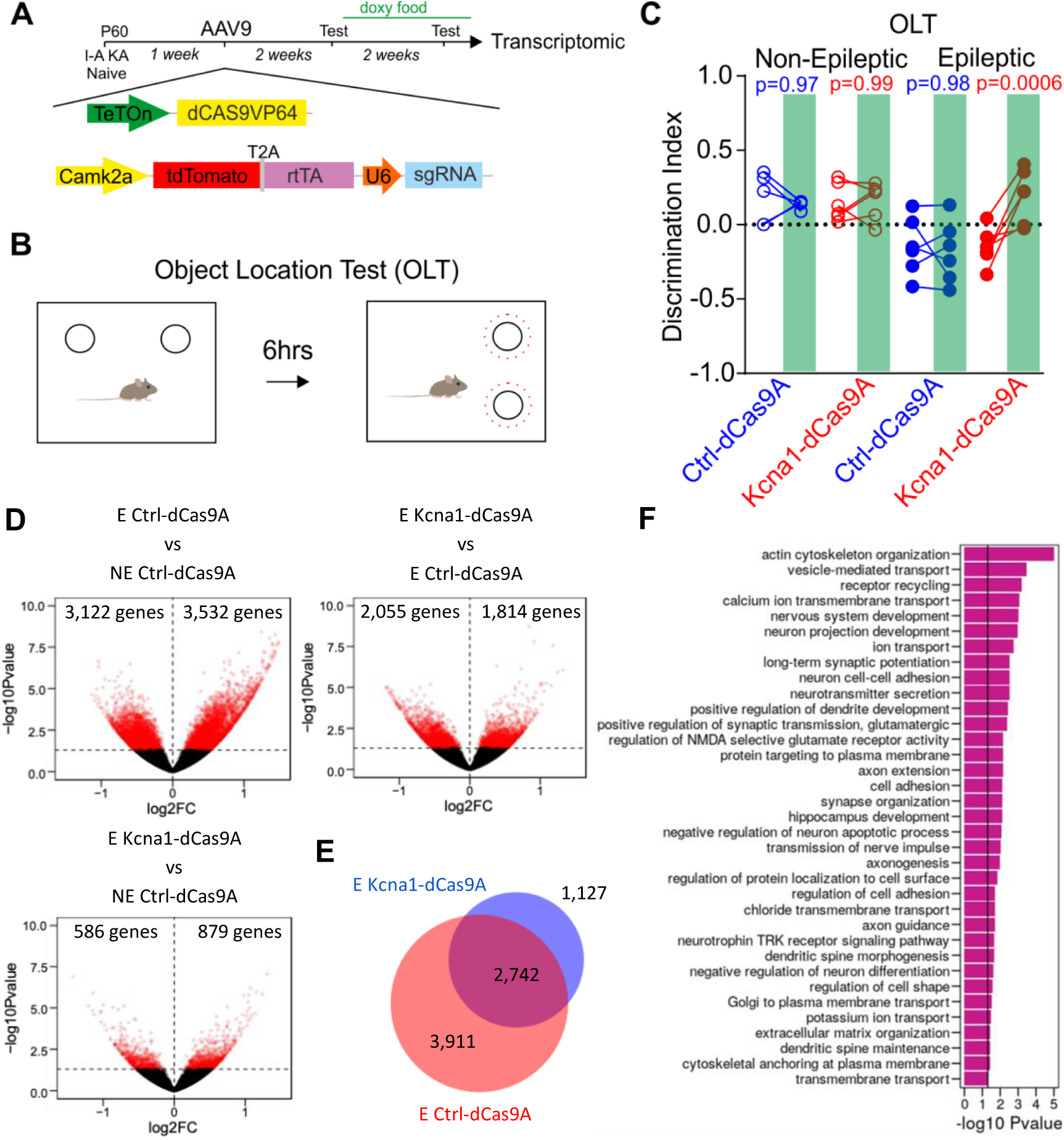
CRISPRa-Kcna1 rescues cognitive deficits and mitigates transcriptomic changes in a mouse model of acquired intractable temporal lobe epilepsy. **A.** Experimental plan for behaviour and transcriptomic analysis. **B.** Graphical representation of the OLT and NORT tests. **C.** Discrimination index for non-epileptic and epileptic animals before and after (green box) doxycycline in mice treated either with Ctrl-dCas9A or Kcna1-dCas9A. Two-way ANOVA followed by Bonferroni multiple comparison test. **D.** Volcano plots showing statistical significance (-log10pvalue) as a function of fold-change in gene expression (log2FC), comparing pairs of datasets as indicated above each plot. Genes showing p<0.05 difference in expression (-log10pvalue>1.3) are highlighted in red. **E.** Venn diagram showing the fraction of differentially regulated genes in epileptic Ctrl-dCas9A-treated mice (E Ctrl-dCas9A) which were rescued in Kcna1-dCas9A-treated mice (E Kcna1-dCas9A).**F.** Histogram displaying representative gene ontology (GO) categories functionally enriched among the 2,742 genes that were differentially expressed in epileptic control mice (E Ctrl-dCas9A) versus treated mice (E Kcna1-dCas9A).

Several studies have documented extensive changes in gene expression in different mouse seizure models (Motti *et al*., 2010; Okamoto *et al*., 2010; Winden *et al*., 2011; Hansen *et al*., 2014; Hawkins *et al*., 2019). We asked if Kcna1-dCas9A is able to rescue transcriptomic changes in the TLE model by performing RNAseq analysis on hippocampi from NE animals injected with Ctrl-dCas9A and E injected either with Ctrl-dCas9A or with Kcna1-dCas9A. A comparison of gene expression between non-epileptic (NE Ctrl-dCas9A) and epileptic animals (E Ctrl-dCas9A) revealed 6654 genes whose expression was altered (Figure 4D, first panel). Kcna1-dCas9A treatment (E Kcna1-dCas9A) (Figure 4E) rescued expression of 2742 of them (Figure 4D, second panel and Figure 4E). Consequently, the transcriptional profile of treated hippocampi (E Kcna1-dCas9A group) was more similar to non-epileptic mice (NE Ctrl-dCas9A group), with only 1465 genes altered (Figure 4D, third bottom panel), than to epileptic Ctrl-dCas9A-treated mice. Sample distribution and K-means clustering along the first two principal components (PC1, PC2) failed to discriminate between NE Ctrl-dCas9A and E Kcna1-dCas9A groups, while the E Ctrl-dCas9A group was distinct (Supplementary Figure 10A). Gene ontology analysis of the 2742 genes rescued by Kcna1-dCas9A treatment (Figure 4F) showed that it counteracted changes implicated in neurodegeneration and apoptosis (GO ID: negative regulation of anti-apoptotic process, hippocampus development, positive regulation of dendrite development, neuronal development with synapse formation, axon extension, axonogenesis, axon guidance). Several genes associated with neuronal activity were upregulated (GO ID: ion transport, calcium transmembrane, chloride transmembrane transport, potassium ion transport). The treatment also re-established expression of genes implicated in synapse function (GO ID: neurotransmitter secretion, synapse organization, dendritic spine maintenance, long term synaptic potentiation, transmission of nerve impulse, receptor recycling) and glutamatergic transmission (GO ID: positive regulation of glutamatergic synaptic transmission, regulation of NMDA selective glutamate receptor activity).

We compared the pattern of gene expression changes in our model with previously published studies (Motti *et al*., 2010; Okamoto *et al*., 2010; Winden *et al*., 2011; Hansen *et al*., 2014; Hawkins *et al*., 2019). This however failed to reveal a consistent signature across heterogeneous models of epilepsy (Supplementary Figure 10B). However, when restricting the comparison to focal kainic acid injection (Motti *et al*., 2010) we identified 388 common deregulated genes, 165 of which were rescued in Kcna1-dCas9A treated mice (Supplementary Figure 10C, D).

## Discussion

Although CRISPR has attracted intense interest as a possible treatment for inherited or acquired genetic disorders, it has, hitherto, received much less attention as a potential tool to treat acquired non-genetic diseases. The overwhelming majority of epilepsy cases, which represent an enormous disease burden, are not thought to be due to single gene mutations but are acquired during life, often secondary to a variety of brain insults such as infections, strokes and injuries (Kwan *et al*., 2011; Tang *et al*., 2017). Here we have shown that CRISPRa can be used to increase endogenous *Kcna1* expression to modulate neuronal activity, to reduce seizure initiation and to rescue behavioural and transcriptomic abnormalities in a mouse model of chronic temporal lobe epilepsy. This approach can potentially be used to regulate the expression of any gene, opening the way to treating many other neurological diseases associated with altered transcription.

At present, the main obstacles to clinical translation of the CRISPR/Cas9 toolbox are absence of long-term data on potential immunogenicity of the bacterial nuclease in humans and possible off-target effects that have not been detected by transcriptomic analysis (Kosicki *et al*., 2018). Although in this study a non-mammalian nuclease has been used, which can evoke an immune response in the primate CNS (Samaranch *et al*., 2014), CNS disorders attributable to antibodies against nuclear neuronal proteins are rare, and rare forms of autoimmune encephalitis generally involve membrane proteins (Platt *et al*., 2017). The doxycycline transactivator protein could also potentially evoke an immune response, although new inducible systems are in consideration for clinical translation (Das *et al*., 2016; Kundert *et al*., 2019). Cas9 has already been delivered with AAV in rodents and has been shown to induce a mild cellular response, but this has not been reported in the brain (Chew *et al*., 2016). CRISPRa is, in principle, less likely to have deleterious off-target effects than CRISPR gene editing because it does not cleave DNA (La Russa and Qi, 2015; Zheng *et al*., 2018), but further research is necessary.

Among distinct advantages of CRISPRa over exogenous gene delivery is the possibility to select one or more sgRNAs to tailor the exact level of gene expression independently by the number of viral copies effectively entered within each neuron. In addition, several sgRNAs could in principle be used in combination to control the transcription of heteromultimeric proteins such as GABA_A_ or NMDA receptors, or of multiple genes in a signalling pathway. Finally, whilst exogenous gene delivery is limited by the viral packaging capacity, CRISPRa can potentially be applied to control the activity of any gene irrespective of its length (Konermann *et al*., 2015).

Although the present study made use of two AAVs to allow inducible activation of CRISPRa and expression of a fluorescent reporter protein, for clinical translation these features would not be required, as it should be possible to package both the dCas9 and the sgRNA in a single AAV to simplify clinical delivery. Further refinements can be considered, such as the use of an inducible promoter to allow the therapy to be switched off (Wykes and Lignani, 2018), which would not be possible with a gene editing strategy.

Importantly, in the present study we observed rescue not only of epilepsy but also of a behavioural co-morbidity and transcriptomic alterations (Figure 4). We tested the effects of epilepsy and dCas9A treatment in two behavioural tests previously studied in epileptic animals (Cho *et al*., 2015), but several other tests, beyond the scope of this study, could be used to further assess the extent of rescue of co-morbidities (Heinrichs and Seyfried, 2006).

Many studies have reported transcriptomic changes associated with epilepsy in different mouse models (Motti *et al*., 2010; Okamoto *et al*., 2010; Winden *et al*., 2011; Hansen *et al*., 2014; Hawkins *et al*., 2019). However, no single gene alteration recurs across all available databases (Figure S9B). This inconsistency can be ascribed to differences in RNA-Seq technologies, epilepsy models, ages of animals, and delay after the insult. Our data add to the available data on gene expression, and provide, to our knowledge, the first evidence that gene therapy for epilepsy can correct dysregulation of a significant subset of genes.

Importantly, the effect of CRISPRa-mediated Kcna1 upregulation on generalised seizures may reflect an overall rescue of the pathology, including co-morbidities and gene expression, and not only an attenuation of acute seizures.

In conclusion, CRISPR-mediated control of gene expression can be successfully exploited to modulate neuronal activity and to mitigate seizures and behavioural co-morbidity in an experimental model of intractable temporal lobe epilepsy.

## Acknowledgments

We thank Jenna Carpenter for the help with neuronal cultures and blinding, and Meena Siddharthana for the analysis of false negative with Pyecog. We are thankful to all members of the DCEE for helpful discussions.

## References

Adli M. The CRISPR tool kit for genome editing and beyond. Nat Commun 2018; 9(1): 1911.

Beaudoin GM, 3rd, Lee SH, Singh D, Yuan Y, Ng YG, Reichardt LF, et al. Culturing pyramidal neurons from the early postnatal mouse hippocampus and cortex. Nat Protoc 2012; 7(9): 1741–54.

Breiman L. Random Forests; 2001.

Carninci P, Sandelin A, Lenhard B, Katayama S, Shimokawa K, Ponjavic J, et al. Genome-wide analysis of mammalian promoter architecture and evolution. Nat Genet 2006; 38(6): 626–35.

Chew WL, Tabebordbar M, Cheng JK, Mali P, Wu EY, Ng AH, et al. A multifunctional AAV-CRISPR-Cas9 and its host response. Nat Methods 2016; 13(10): 868–74.

Cho KO, Lybrand ZR, Ito N, Brulet R, Tafacory F, Zhang L, et al. Aberrant hippocampal neurogenesis contributes to epilepsy and associated cognitive decline. Nat Commun 2015; 6: 6606.

Colasante G, Lignani G, Rubio A, Medrihan L, Yekhlef L, Sessa A, et al. Rapid Conversion of Fibroblasts into Functional Forebrain GABAergic Interneurons by Direct Genetic Reprogramming. Cell Stem Cell 2015; 17(6): 719–34.

Das AT, Tenenbaum L, Berkhout B. Tet-On Systems For Doxycycline-inducible Gene Expression. Curr Gene Ther 2016; 16(3): 156–67.

Dobin A, Davis CA, Schlesinger F, Drenkow J, Zaleski C, Jha S, et al. STAR: ultrafast universal RNA-seq aligner. Bioinformatics 2013; 29(1): 15–21.

Dominguez AA, Lim WA, Qi LS. Beyond editing: repurposing CRISPR-Cas9 for precision genome regulation and interrogation. Nat Rev Mol Cell Biol 2016; 17(1): 5–15.

Hansen KF, Sakamoto K, Pelz C, Impey S, Obrietan K. Profiling status epilepticus-induced changes in hippocampal RNA expression using high-throughput RNA sequencing. Sci Rep 2014; 4: 6930.

Hawkins NA, Calhoun JD, Huffman AM, Kearney JA. Gene expression profiling in a mouse model of Dravet syndrome. Exp Neurol 2019; 311: 247–56.

Heeroma JH, Henneberger C, Rajakulendran S, Hanna MG, Schorge S, Kullmann DM. Episodic ataxia type 1 mutations differentially affect neuronal excitability and transmitter release. Dis Model Mech 2009; 2(11-12): 612–9.

Heidenreich M, Zhang F. Applications of CRISPR-Cas systems in neuroscience. Nat Rev Neurosci 2016; 17(1): 36–44.

Heinrichs SC, Seyfried TN. Behavioral seizure correlates in animal models of epilepsy: a road map for assay selection, data interpretation, and the search for causal mechanisms. Epilepsy Behav 2006; 8(1): 5–38.

Jimenez-Mateos EM, Engel T, Merino-Serrais P, McKiernan RC, Tanaka K, Mouri G, et al. Silencing microRNA-134 produces neuroprotective and prolonged seizure-suppressive effects. Nat Med 2012; 18(7): 1087–94.

Katzel D, Nicholson E, Schorge S, Walker MC, Kullmann DM. Chemical-genetic attenuation of focal neocortical seizures. Nat Commun 2014; 5: 3847.

Konermann S, Brigham MD, Trevino AE, Joung J, Abudayyeh OO, Barcena C, et al. Genome-scale transcriptional activation by an engineered CRISPR-Cas9 complex. Nature 2015; 517(7536): 583–8.

Kosicki M, Tomberg K, Bradley A. Repair of double-strand breaks induced by CRISPR-Cas9 leads to large deletions and complex rearrangements. Nat Biotechnol 2018.

Krook-Magnuson E, Armstrong C, Oijala M, Soltesz I. On-demand optogenetic control of spontaneous seizures in temporal lobe epilepsy. Nat Commun 2013; 4: 1376.

Kullmann DM, Schorge S, Walker MC, Wykes RC. Gene therapy in epilepsy-is it time for clinical trials? Nat Rev Neurol 2014; 10(5): 300–4.

Kundert K, Lucas JE, Watters KE, Fellmann C, Ng AH, Heineike BM, et al. Controlling CRISPR-Cas9 with ligand-activated and ligand-deactivated sgRNAs. Nat Commun 2019; 10(1): 2127.

Kwan P, Schachter SC, Brodie MJ. Drug-resistant epilepsy. N Engl J Med 2011; 365(10): 919–26.

La Russa MF, Qi LS. The New State of the Art: Cas9 for Gene Activation and Repression. Mol Cell Biol 2015; 35(22): 3800–9.

Leger M, Quiedeville A, Bouet V, Haelewyn B, Boulouard M, Schumann-Bard P, et al. Object recognition test in mice. Nat Protoc 2013; 8(12): 2531–7.

Liao HK, Hatanaka F, Araoka T, Reddy P, Wu MZ, Sui Y, et al. In Vivo Target Gene Activation via CRISPR/Cas9-Mediated Trans-epigenetic Modulation. Cell 2017; 171(7): 1495–507 e15.

Lieb A, Qiu Y, Dixon CL, Heller JP, Walker MC, Schorge S, et al. Biochemical autoregulatory gene therapy for focal epilepsy. Nat Med 2018.

Lopes G, Bonacchi N, Frazao J, Neto JP, Atallah BV, Soares S, et al. Bonsai: an event-based framework for processing and controlling data streams. Front Neuroinform 2015; 9: 7.

Love MI, Huber W, Anders S. Moderated estimation of fold change and dispersion for RNA-seq data with DESeq2. Genome Biol 2014; 15(12): 550.

Magloire V, Cornford J, Lieb A, Kullmann DM, Pavlov I. KCC2 overexpression prevents the paradoxical seizure-promoting action of somatic inhibition. Nat Commun 2019; 10(1): 1225.

Morabito G, Giannelli SG, Ordazzo G, Bido S, Castoldi V, Indrigo M, et al. AAV-PHP.B-Mediated Global-Scale Expression in the Mouse Nervous System Enables GBA1 Gene Therapy for Wide Protection from Synucleinopathy. Mol Ther 2017; 25(12): 2727–42.

Morris G, Leite M, Kullmann DM, Pavlov I, Schorge S, Lignani G. Activity Clamp Provides Insights into Paradoxical Effects of the Anti-Seizure Drug Carbamazepine. J Neurosci 2017; 37(22): 5484–95.

Motti D, Le Duigou C, Eugene E, Chemaly N, Wittner L, Lazarevic D, et al. Gene expression analysis of the emergence of epileptiform activity after focal injection of kainic acid into mouse hippocampus. Eur J Neurosci 2010; 32(8): 1364–79.

Okamoto OK, Janjoppi L, Bonone FM, Pansani AP, da Silva AV, Scorza FA, et al. Whole transcriptome analysis of the hippocampus: toward a molecular portrait of epileptogenesis. BMC Genomics 2010; 11: 230.

Pinatel D, Hivert B, Saint-Martin M, Noraz N, Savvaki M, Karagogeos D, et al. The Kv1-associated molecules TAG-1 and Caspr2 are selectively targeted to the axon initial segment in hippocampal neurons. J Cell Sci 2017; 130(13): 2209–20.

Platt MP, Agalliu D, Cutforth T. Hello from the Other Side: How Autoantibodies Circumvent the Blood-Brain Barrier in Autoimmune Encephalitis. Front Immunol 2017; 8: 442.

Pozzi D, Lignani G, Ferrea E, Contestabile A, Paonessa F, D’Alessandro R, et al. REST/NRSF-mediated intrinsic homeostasis protects neuronal networks from hyperexcitability. EMBO J 2013; 32(22): 2994–3007.

Samaranch L, Sebastian WS, Kells AP, Salegio EA, Heller G, Bringas JR, et al. AAV9-mediated expression of a non-self protein in nonhuman primate central nervous system triggers widespread neuroinflammation driven by antigen-presenting cell transduction. Mol Ther 2014; 22(2): 329–37.

Simonato M. Gene therapy for epilepsy. Epilepsy Behav 2014; 38: 125–30.

Snowball A, Chabrol E, Wykes RC, Shekh-Ahmad T, Cornford JH, Lieb A, et al. Epilepsy Gene Therapy Using an Engineered Potassium Channel. J Neurosci 2019; 39(16): 3159–69.

Suzuki K, Tsunekawa Y, Hernandez-Benitez R, Wu J, Zhu J, Kim EJ, et al. In vivo genome editing via CRISPR/Cas9 mediated homology-independent targeted integration. Nature 2016; 540(7631): 144–9.

Tang F, Hartz AMS, Bauer B. Drug-Resistant Epilepsy: Multiple Hypotheses, Few Answers. Front Neurol 2017; 8: 301.

Thorvaldsdottir H, Robinson JT, Mesirov JP. Integrative Genomics Viewer (IGV): high-performance genomics data visualization and exploration. Brief Bioinform 2013; 14(2): 178–92.

Vivekananda U, Novak P, Bello OD, Korchev YE, Krishnakumar SS, Volynski KE, et al. Kv1.1 channelopathy abolishes presynaptic spike width modulation by subthreshold somatic depolarization. Proc Natl Acad Sci U S A 2017; 114(9): 2395–400.

Watanabe I, Wang HG, Sutachan JJ, Zhu J, Recio-Pinto E, Thornhill WB. Glycosylation affects rat Kv1.1 potassium channel gating by a combined surface potential and cooperative subunit interaction mechanism. J Physiol 2003; 550(Pt 1): 51–66.

Winden KD, Karsten SL, Bragin A, Kudo LC, Gehman L, Ruidera J, et al. A systems level, functional genomics analysis of chronic epilepsy. PLoS One 2011; 6(6): e20763.

Wykes RC, Heeroma JH, Mantoan L, Zheng K, MacDonald DC, Deisseroth K, et al. Optogenetic and potassium channel gene therapy in a rodent model of focal neocortical epilepsy. Sci Transl Med 2012; 4(161): 161ra52.

Wykes RC, Kullmann DM, Pavlov I, Magloire V. Optogenetic approaches to treat epilepsy. J Neurosci Methods 2016; 260: 215–20.

Wykes RC, Lignani G. Gene therapy and editing: Novel potential treatments for neuronal channelopathies. Neuropharmacology 2018; 132: 108–17.

Zheng Y, Shen W, Zhang J, Yang B, Liu YN, Qi H, et al. CRISPR interference-based specific and efficient gene inactivation in the brain. Nat Neurosci 2018; 21(3): 447–54.

