## Supplementary Materials for "*In vivo* CRISPRa decreases seizures and rescues cognitive deficits in a rodent model of epilepsy"

**Table 1. sgRNA and primer sequences**

| **Name** | **Sequence** |
|---|---|
| sglacZ | TGCGAATACGCCCACGCGAT |
| Sg4 | CTTCGCACCACCGCGGCTCG CGG |
| Sg14 | GGGGCCTCTAGAAGGATCCC AGG |
| Sg19 | AGTCAATGATCACATCCTCC TGG |
| Sg30 | ACCCCAGCATTCCTTTCAGG GGG |
| 18s_F | GGTGAAATTCTTGGACCGGC |
| 18s_R | GACTTTGGTTTCCCGGAAGC |
| Kcna1_F | TTCAGACTCTCCGCCGACTC |
| Kcna1_R | CAGGCCTGTCACCCACTTTG |
| dCas9-VP64_F | TCCTTTTTGGTGGAGGAGGA |
| dCas9-VP64_R | TCAACCGCAAGTCAGCCTTA |
| Pde4b_F | TCAGCCAGGTCTAATCTGCCA |
| Pde4b_R | ACCGCCATCACACTCCTACT |
| Mylpf_F | CTCACCTCCCCGAATGTCCT |
| Mylpf_R | GGGTCCCCTATTCCTGGTCC |

| | |
|---|---|
| Efcab4a_F | CCAGCATTCCACCATTCCCC |
| Efcab4a_R | GTCGACTGAGCTGCTCTTGG |
| Nudcd2_F | CCGAGCCCTAAAATTCACGGT |
| Nudcd2_R | ACAGTTTCCCCTTGCCACAC |
| Gc_F | AGGAGGTGCTGCAAGACTCT |
| Gc_R | CCTTCTCATAGTCTCGGCCTCT |
| Vps16_F | ACACTGCGAACTGGAATCCAC |
| Vps16_R | GCAATCCTTGAGCTCCTCCTTC |

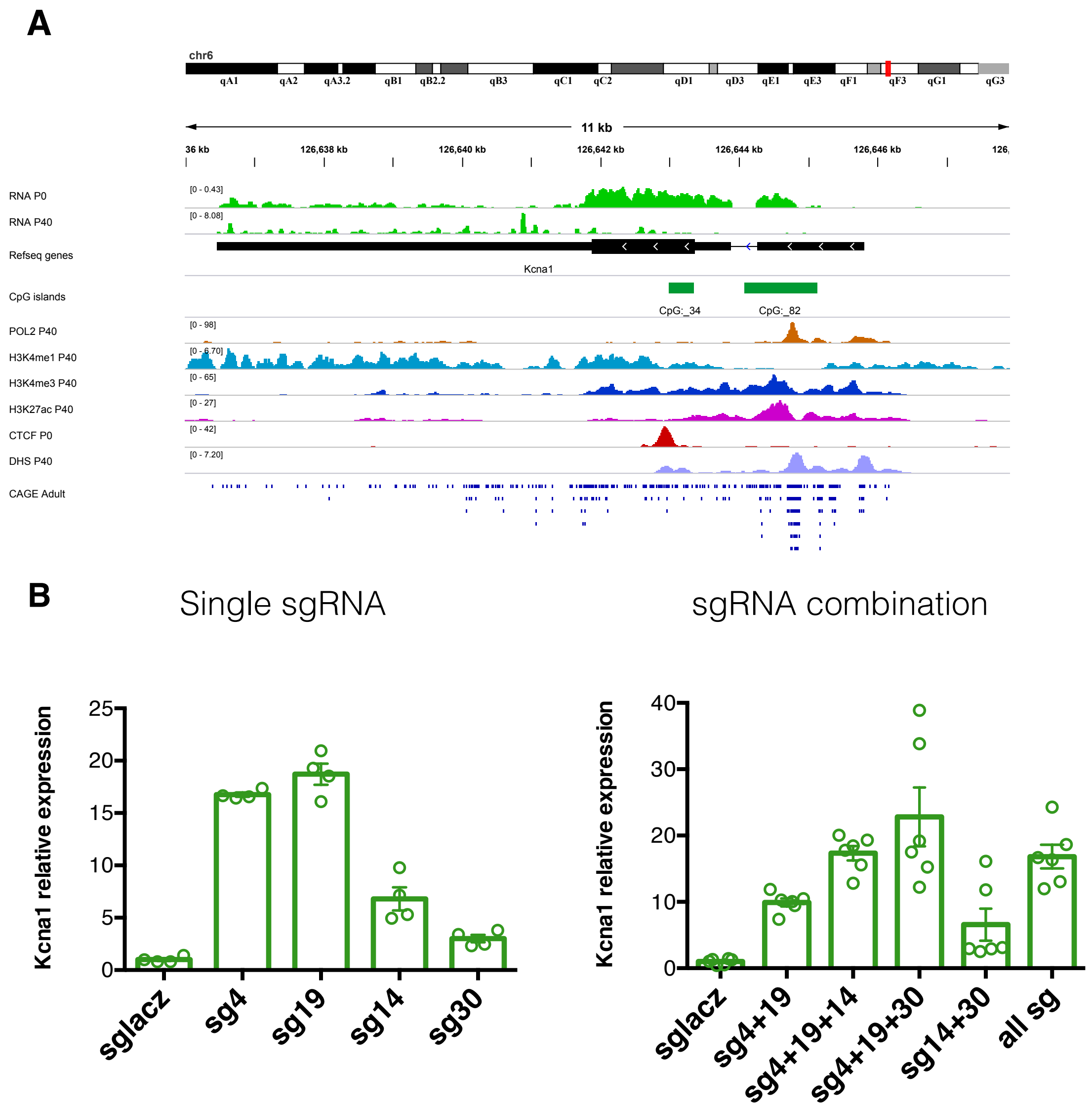


**Figure S1: Bioinformatics analysis for prediction of *Kcna1* gene promoter and sgRNA screening for stimulating *Kcna1* gene expression with CRISPRa in P19 cells. A**. Alignment of *Kcna1* gene reference sequence with RNA-seq of mouse brain at P0 and P40, ChIP-seq at P40 for POL2 (RNA polymerase II), H3K4me1 (mono-methylation of lysine 4 on histone H3), H3K4me3 (tri-methylation of lysine 4 on the histone H3) and H3K27ac (acetylation of lysine 27 on histone H3), CTCF (factor that binds the CCCTC ) and DNase-seq (DHS, DNase I Hyper Sensitivity mapping) and CAGE- seq (Cap Analysis of Gene Expression-sequencing) profiles. The enrichment of markers associated with transcriptional activation in the regions upstream to the first exon of the gene highlight the presence of a TSS and allows to localize a promoter region in the 200 bp upstream. **B**. RT-qPCRs for *Kcna1* mRNA levels on RNA extracted from P19 cells lipofected with dCas9VP160-T2A-Puro$^R$ together with sgRNAs targeting Kcna1 gene promoter. Data are normalized on the 18S rRNA and relative to control sgLacZ cells.

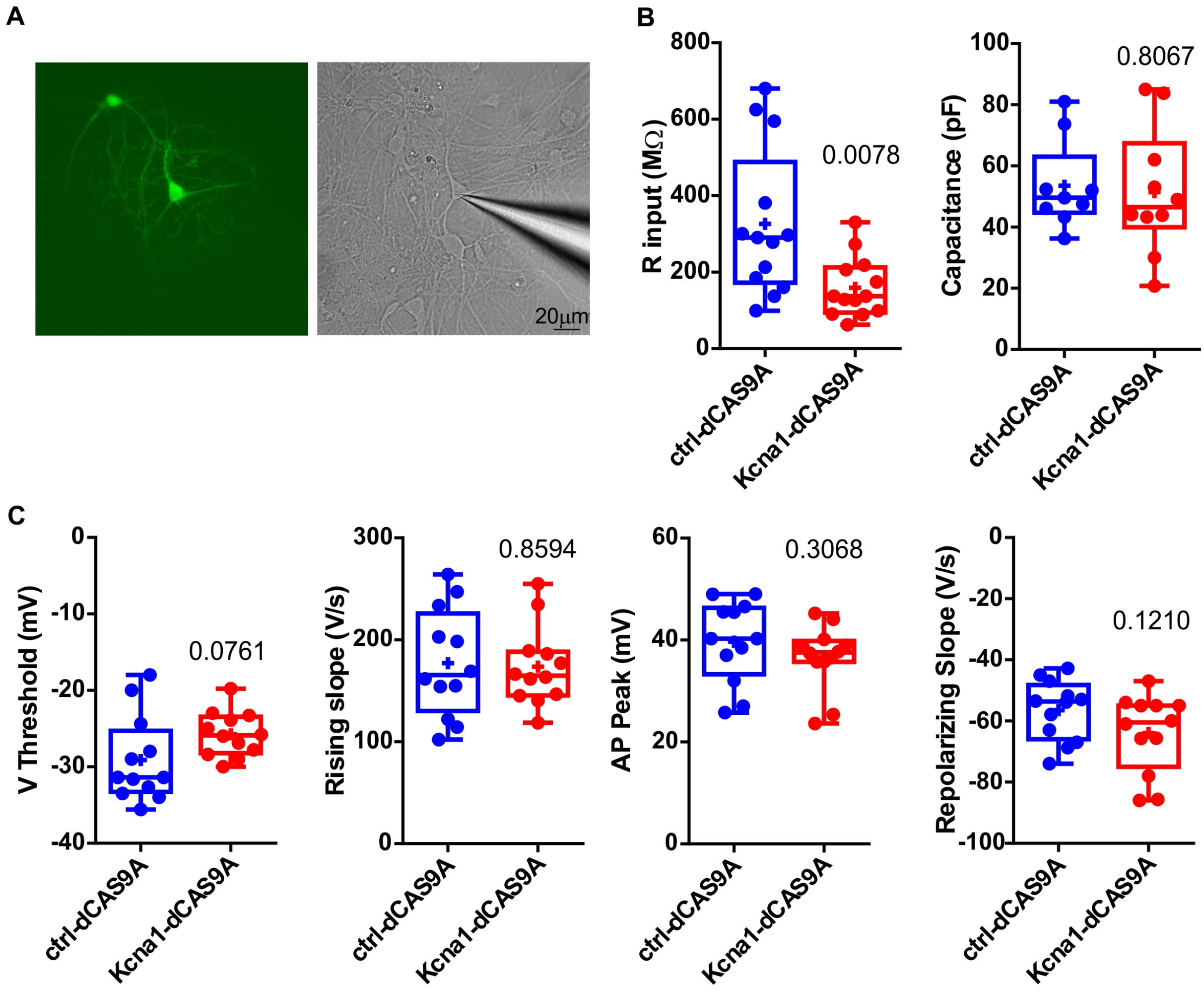


**Figure S2: Electrophysiology recordings from cultured neurons. Neuronal and AP parameters. A.** representative picture of a patched transduced neuron in vitro (Green= EGFP). **B, C.** Cell (B) and AP (C) parameters of recorded cells. Student's t test. Only the change in input resistance was significant for Ctrl-dCas9A compared to Kcna1-dCas9A neurons.

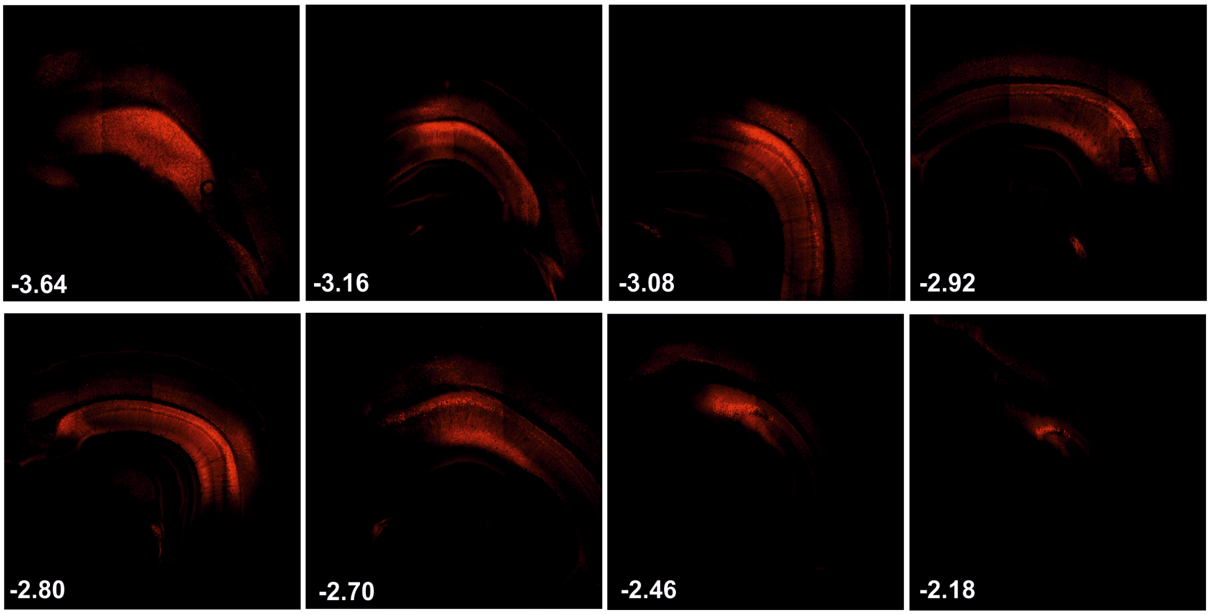


**Figure S3: tdTomato expression in slices from Camk2a/CRE mouse.** Representative images of slices from a Camk2a/CRE mouse injected with Ctrl-dCas9A. Native tdTomato (non-immunofluorescence) was present in the floxed rtTa-t2a-tdTomato cassette driven by Syn promoter. Coordinates are from Bregma.

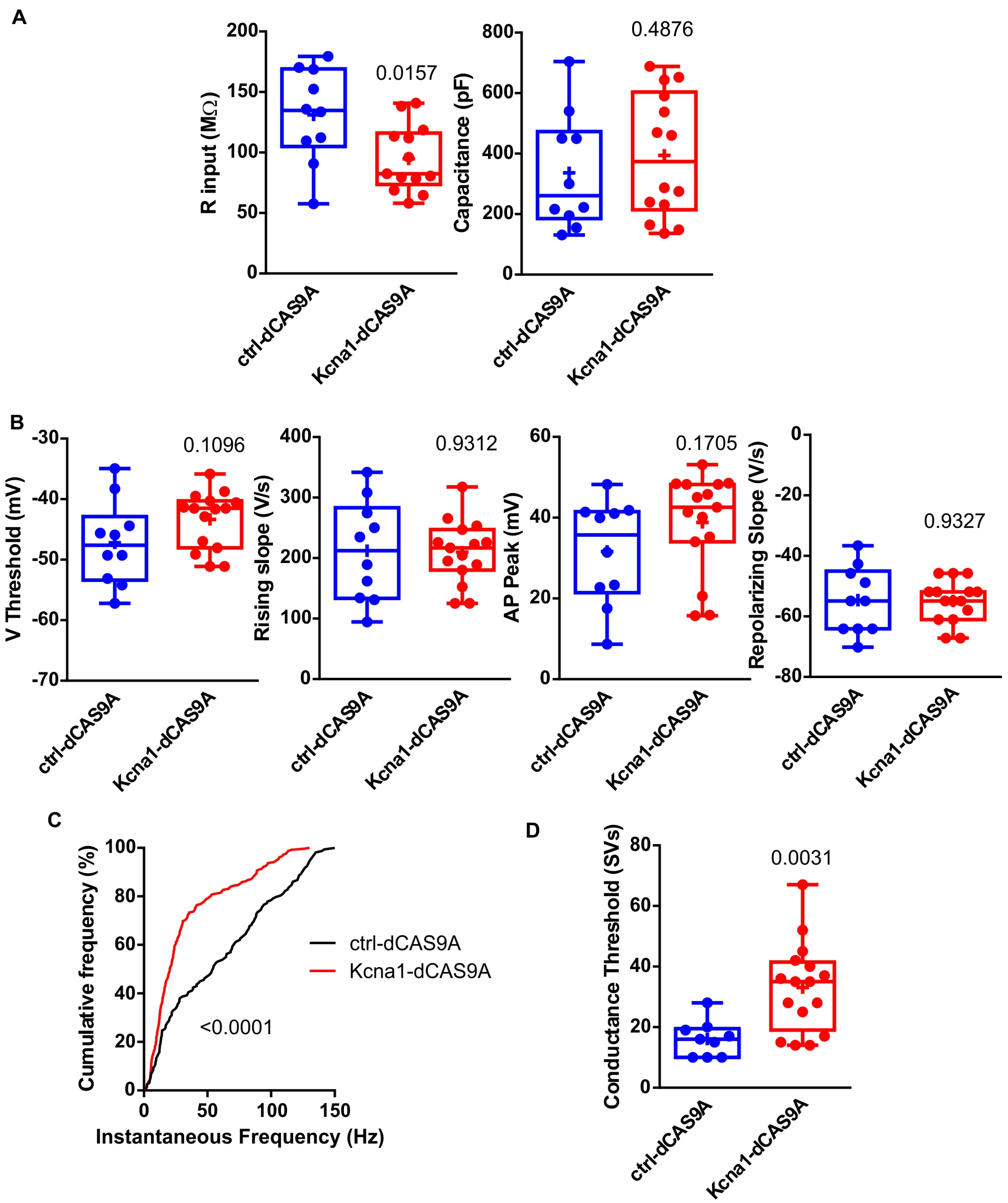


**Figure S4:** ***Ex vivo*** **electrophysiological recordings in acute slices from Camk2a/CRE mice. A, B.** Cell (B) and AP parameters. Student's t test. Only the input resistance was significant for Ctrl-dCs9A compared to Kcna1-dCs9A neurons. **C.** Cumulative analysis of instantaneous frequency for the first 2 APs in each current step for all neurons transduced either with Ctrl-dCas9A or Kcna1-dCs9A. Mann-Whitney non-parametric test. **D.** Conductance threshold calculated as the first AP elicited with steps of simulated single AMPA miniature events (Morris *et al.*, 2017).

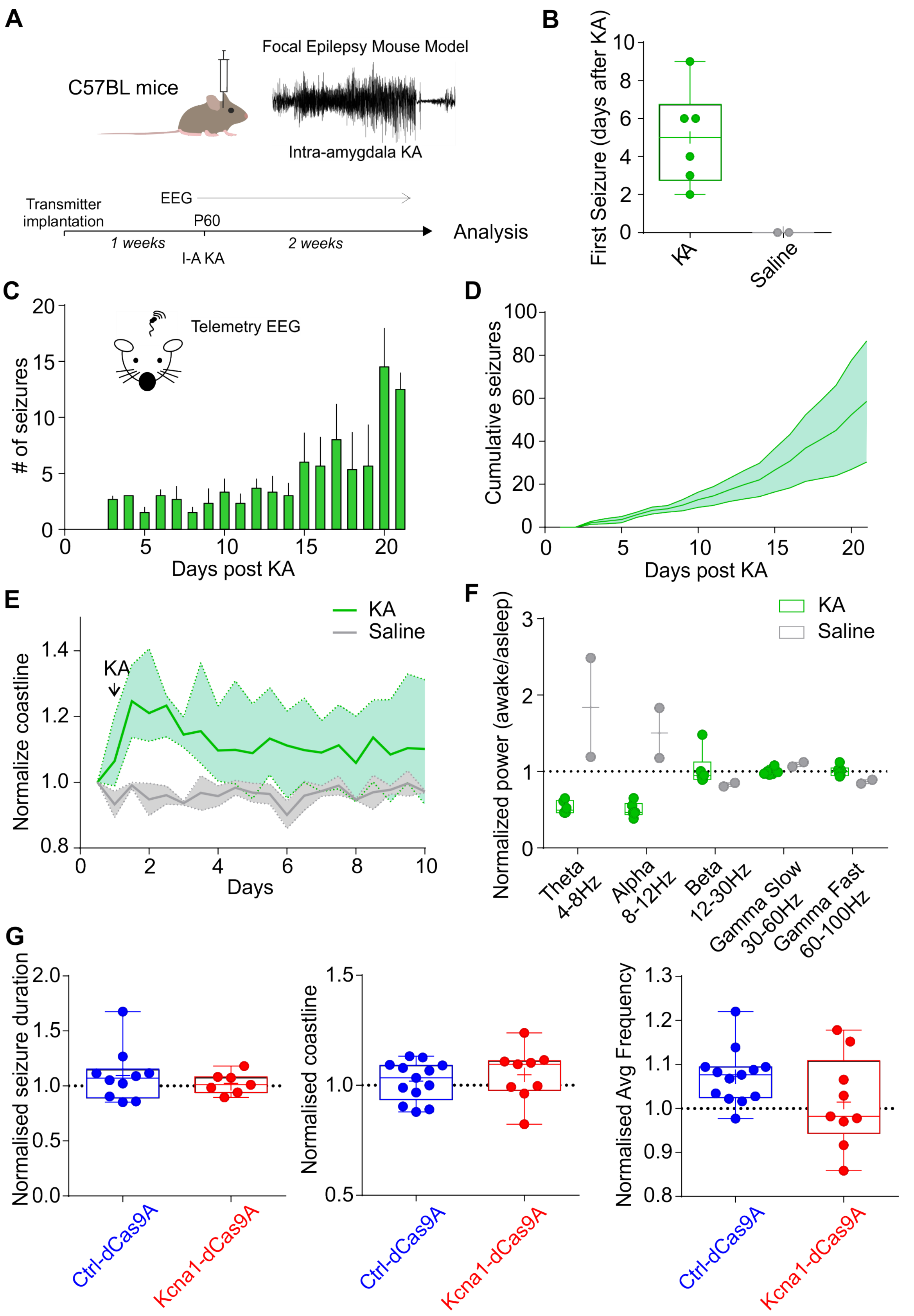


**Figure S5: Intra-amygdala kainic acid model of temporal lobe epilepsy. A.** Experimental plan for inducing chronic epilepsy injecting KA in the right amygdala to induce status epilepticus (SE). **B.** Days after KA before a stage 5 generalized tonic-clonic seizure in animals injected with KA or saline in amygdala. Average first seizure occurred 5 days after KA. **C, D.** Number of seizures (C) and cumulative number of seizures (D) in the first 3 weeks after KA injection (n=6). **E.** Coastline in the first 10 days after SE normalized to 12hrs EEG recordings before SE for animals injected with KA (n=6) or saline (n=2). Data points are binned every 12hrs. **F.** Power during wakefulness (defined as the 12hr dark period) as a ratio of power during sleep (defined as the 12hr light period) for the theta, alpha, beta, slow and fast gamma bands. **G.** Seizure duration, coastline and power in the different bands recorded after doxycycline, normalized by values before doxycycline, did not differ between groups. The normalised averaged frequency was calculated as the sum over all bands: (Power in band /sum of all power) * middle of the power band.

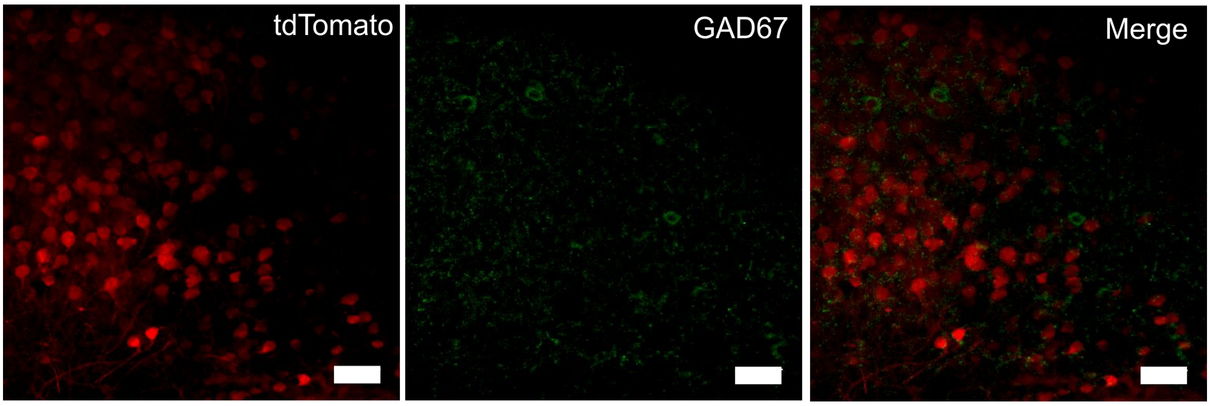


**Figure S6: Immunohistochemistry of epileptic brains after the recordings.** Representative image of a Ctrl-dCas9A injected mouse showing no co-localization between tdTomato, driven by Camk2a promoter, and GAD67, a marker for inhibitory neurons (scale bar: 50µm). No co-localisation between Camk2a-driven tdTomato and GAD67 was detected in any of the 4 brains analysed after recordings.

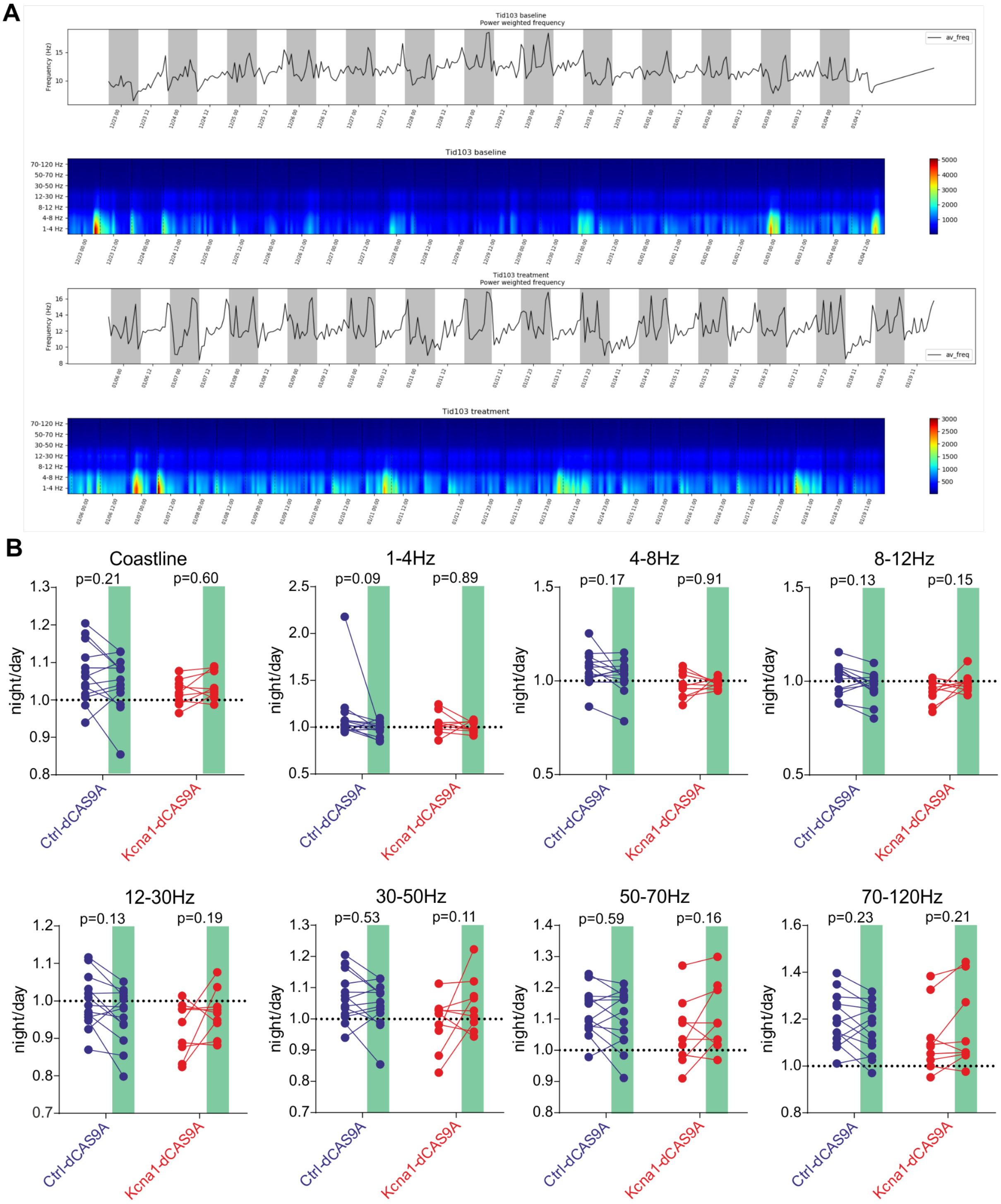


**Figure S7. CRISPRa-Kcna1 did not differentially affect EEG power during night or day, in epileptic mice. A.** Representative spectrogram of total frequency and of individual frequencies before (top panels) and after doxycycline (bottom panels) **B.** Quantification of coastline and EEG power at different frequencies as ratio between night and day. No effect of doxycycline was observed in control- and Kcna1-dCas9A treated animals. Two-way ANOVA followed by Bonferroni multiple comparison test.

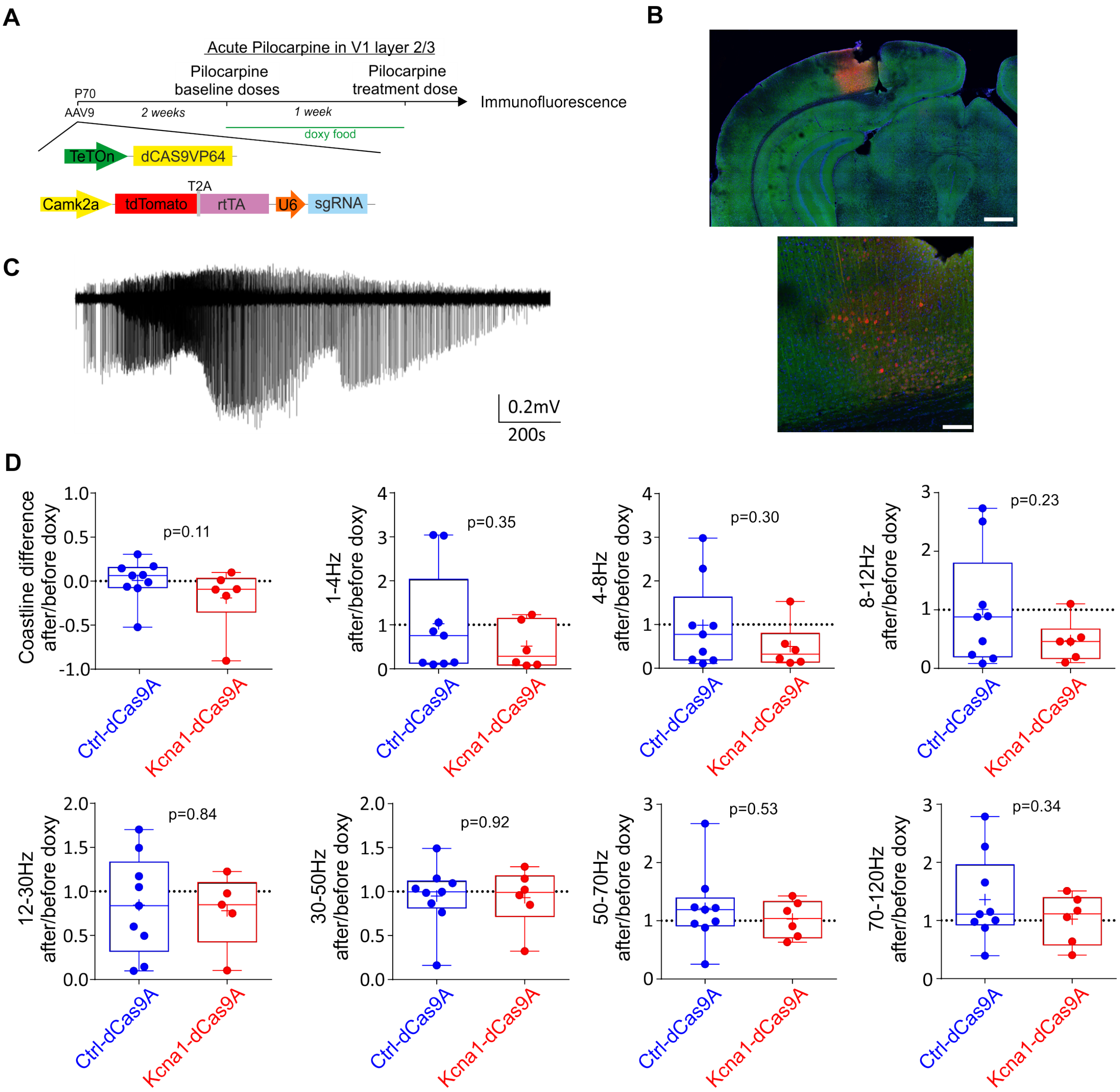


**Figure S8: Acute pilocarpine injection. A.** Graphical experimental design (see Material and methods). **B.** Representative coronal image from a Ctrl-dCas9A injected mouse at the site of injection, with transduced neurons in red, MAP2 in green and DAPI in blue. Scale bar top: 0.5mm; bottom: 0.1mm. **C.** Representative acute seizure induced by pilocarpine. **D.** Seizure parameters after and before doxycycline administration. Coastline measured either before or after doxycycline was normalized by a period before pilocarpine injection, and is shown as the difference (after – before doxycycline). Student's t test.

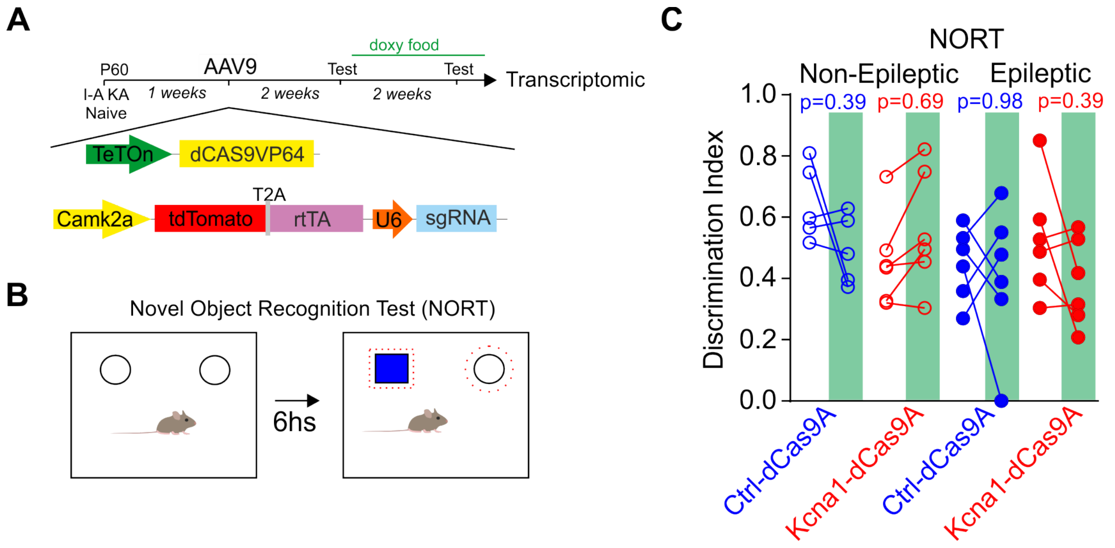


**Figure S9: CRISPRa-Kcna1 doesn't change behavior performance in a test not related with hippocampus. A**. Experimental plan for behaviour analysis. **B**. Graphical representation of NORT tests. **C**. Discrimination index for non-epileptic and epileptic animals before and after (green box) doxycycline in mice treated either with Ctrl-dCas9A or Kcna1-dCas9A. Two-way ANOVA followed by Bonferroni multiple comparison test.

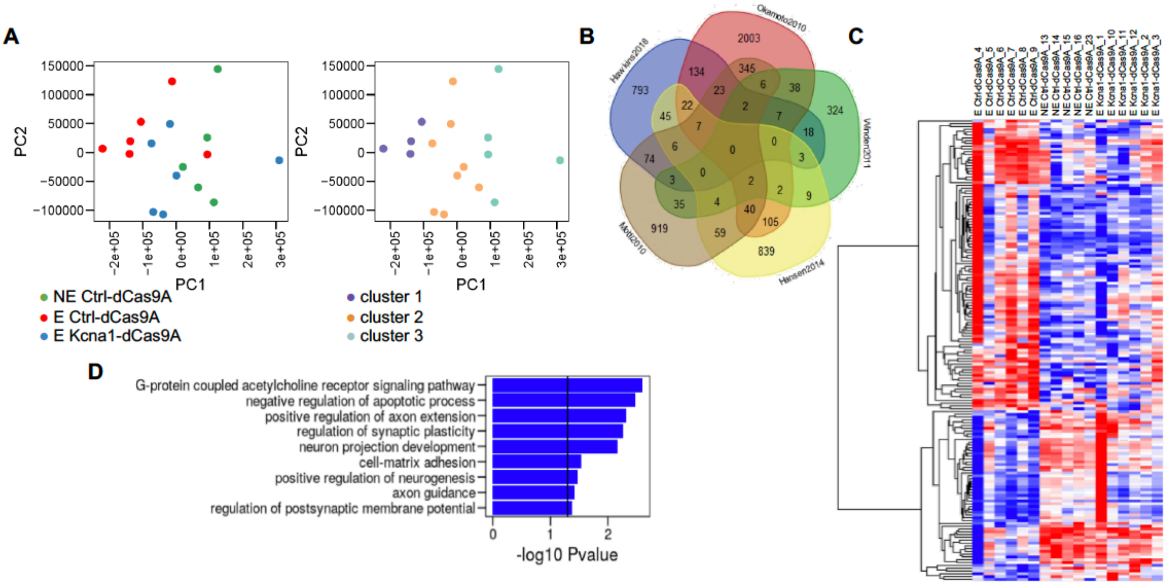


**Figure S10**: **Changes in gene transcription in the epilepsy model**. **A**. Sample distribution along principal component one (PC1) and two (PC2) of whole transcriptome Principal Component Analysis (PCA), color-coded according to sample group or K-Means machine learning clustering. **B**. Venn diagram displaying the genes reported as differentially expressed in genetic (Scn1a +/-) and pharmacologically induced (Kainate, Pilocarpine) murine epilepsy models, with the source literature indicated.

**C**. Gene expression heatmap showing 165 dysregulated genes both in kainate-treated epileptic mice (Motti et al., 2010) and this study, and partial rescue with the Kcna1-dCas9A treatment.

**C**. Histogram displaying representative gene ontology (GO) categories that are functionally enriched among the aforementioned 165 genes.

**Video S1:** Representative video of SE in a C57BL/6J mouse (2minutes from a 40minutes SE).

**Video S2:** Representative video of a Stage 5 generalized tonic-clonic seizure in a C57Bl/6J mouse 4 weeks after KA injection and transduced with ctrl-dCAS9A.
